# A visual pathway for skylight polarization processing in *Drosophila*

**DOI:** 10.1101/2020.09.10.291955

**Authors:** Ben J. Hardcastle, Jaison J. Omoto, Pratyush Kandimalla, Bao-Chau M. Nguyen, Mehmet F. Keleş, Natalie K. Boyd, Volker Hartenstein, Mark A. Frye

## Abstract

Many insects use patterns of polarized light in the sky to orient and navigate. Here we functionally characterize neural circuitry in the fruit fly, *Drosophila melanogaster*, that conveys polarized light signals from the eye to the central complex, a brain region essential for the fly’s sense of direction. Neurons tuned to the angle of polarization of ultraviolet light are found throughout the anterior visual pathway, connecting the optic lobes with the central complex via the anterior optic tubercle and bulb, in a homologous organization to the ‘sky compass’ pathways described in other insects. We detail how a consistent, map-like organization of neural tunings in the peripheral visual system is transformed into a reduced representation suited to flexible processing in the central brain. This study identifies computational motifs of the transformation, enabling mechanistic comparisons of multisensory integration and central processing for navigation in the brains of insects.

## INTRODUCTION

A critical challenge of active locomotion is knowing the right way to go. Sensorimotor reflexes can influence momentary changes in direction to hold a course or to avoid looming threats, but goal-directed behaviors, such as returning to a previous location from unfamiliar surroundings, require additional information and processing (Braitenberg, 1986; Gomez-Marin et al., 2010). External sensory cues must be transformed into an internal representation of position and orientation within the environment, which can also be modified by past experience (Collett and Collett, 2002). In Dipteran flies, as in other invertebrates, a collection of neuropils known as the central complex (CX) is believed to coordinate such behaviors and plays a role in spatial memory, object memory, and action selection (Giraldo et al., 2018; Neuser et al., 2008; Ofstad et al., 2011; Strausfeld and Hirth, 2013), in addition to homeostatic processes including hunger and sleep (Donlea et al., 2014; Dus et al., 2013; Liu et al., 2016).

Recent studies in *Drosophila* have revealed that activity in a network of CX neurons encodes and maintains a representation of the animal’s angular heading relative to its environment (Kim et al., 2017; Seelig and Jayaraman, 2015), with similarity to head-direction cells in vertebrates (Taube et al., 1990). This neural representation of heading can be updated by internal, proprioceptive estimates of self-motion during locomotion, and by external cues, such as moving visual patterns and directional airflow (Fisher et al., 2019; Green et al., 2017; Kim et al., 2019; Okubo et al., 2020; Shiozaki et al., 2020). In other insects, including locusts, crickets, bees, butterflies, and beetles, the functional organization of the CX has frequently been studied in the context of navigation via celestial cues, particularly polarized light (Heinze, 2014). The nearly ever-present pattern of polarization in the sky, formed by scattering of light in the atmosphere, offers an indicator of orientation to organisms able to detect and interpret it, and may be more stable than terrestrial landmarks (Cronin and Marshall, 2011; Dacke et al., 2003; v. Frisch, 1949; Horváth and Varju, 2004; Mappes and Homberg, 2004; Wehner and Müller, 2006). In these non-Dipteran insects, a multimodal neural circuit transmits polarization signals from the eyes to the central complex (Heinze, 2013; Heinze and Reppert, 2011; Homberg et al., 2011; el Jundi et al., 2014, 2015; Pfeiffer et al., 2005). This circuit is known as the ‘sky compass’ pathway for its proposed role in processing skylight polarization patterns and information about the position of the sun to bestow an animal with a sense of direction. In *Drosophila*, the anterior visual pathway (AVP), which comprises neurons connecting the medulla, anterior optic tubercle, bulb, and ellipsoid body, has been postulated to represent the homologue of the sky compass pathway (Omoto et al., 2017; Timaeus et al., 2017; Warren et al., 2019). Visual processing in the AVP appears to be segregated into three topographically-organized, parallel streams, of which two have been shown to encode distinct small-field, unpolarized stimuli (Omoto et al., 2017; Seelig and Jayaraman, 2013; Shiozaki and Kazama, 2017; Sun et al., 2017). The neurons involved in polarization processing in *Drosophila* have not been identified beyond peripheral circuits of the dorsal rim area, a specialized region of the eye for detecting skylight polarization (Fortini and Rubin, 1991; Wada, 1974; Weir and Dickinson, 2015; Weir et al., 2016; Wernet et al., 2012; Wolf et al., 1980).

A detailed mapping of the relevant polarization-sensitive neurons would allow the exquisite genetic tools and connectomic studies available in *Drosophila* (Scheffer et al., 2020) to be leveraged to understand the workings of the CX and its integration of multiple sensory modalities. Behavioral experiments have demonstrated that *Drosophila* orient relative to polarization patterns while walking and in tethered-flight (Mathejczyk and Wernet, 2019; Stephens et al., 1953; Warren et al., 2018; Weir and Dickinson, 2011; Wernet et al., 2012; Wolf et al., 1980). A comparative approach would therefore provide insight into the processing strategies employed across taxa as well as species-specific adaptations (Honkanen et al., 2019). Furthermore, it may be possible to reconcile the existing evidence of a common, fixed representation of polarization patterns in the CX of non-Dipteran insects (Heinze and Homberg, 2007; Heinze and Reppert, 2011; Stone et al., 2017) with the emerging model of a flexible representation of both visual information and heading direction in the *Drosophila* CX (Fisher et al., 2019; Kim et al., 2017, 2019; Seelig and Jayaraman, 2015; Turner-Evans et al., 2020). Alternatively, fundamental differences in the organization and processing of polarized light signals between species may reflect specialized navigational requirements.

Here, we set out to test the hypothesis that the anterior visual pathway conveys polarized light signals from the eye to the central complex in *Drosophila*. We used neurogenetic tracing techniques and in vivo calcium imaging to characterize the organization of the neurons at each stage and their coding and transformation of visual features. We show that parallel circuitry in the medulla conducts polarization signals from photoreceptors in the dorsal rim area to a stereotyped domain of the anterior optic tubercle. From there, a postsynaptic population of neurons projecting to the anterior bulb relays polarization signals to ring neurons of the ellipsoid body, and in turn, the ‘compass neurons’ of the central complex. The superior bulb multiplexes polarized and unpolarized light signals, while the inferior bulb does not appear to be involved in polarization processing. Finally, we examine population responses in the central complex and find hallmarks of a flexible encoding of a single angle of polarization which could be used to direct motor output for navigation behavior.

## RESULTS

In flies, the pair of inner photoreceptors in each ommatidium, R7/R8, are involved in the detection of color and linear polarization of light (Hardie, 1984). Within a narrow strip of skyward-facing ommatidia in each eye, known as the dorsal rim area (DRA), each R7/R8 pair is sensitive to a different angle of polarization (AoP, also referred to as the e-vector orientation), organized in a ‘polarotopic’ fashion (Fig. 1A). This specialized array of polarization detectors covers the complete 180° range of orientations and, with a peak spectral sensitivity to UV light, is well-suited to sensing the patterns of polarized light in the sky (Feiler et al., 1992; Salcedo et al., 1999; Sharkey et al., 2020; Weir et al., 2016). A previous characterization of DRA R7/R8 in *Drosophila* established the spatial organization of their tunings, and their visual response properties (Weir et al., 2016). Here, we followed the pathway for skylight polarization signals from the eye and investigated direct downstream targets of DRA R7/R8s at their axon terminals in the second optic neuropil, the medulla (ME).

**Figure 1:**
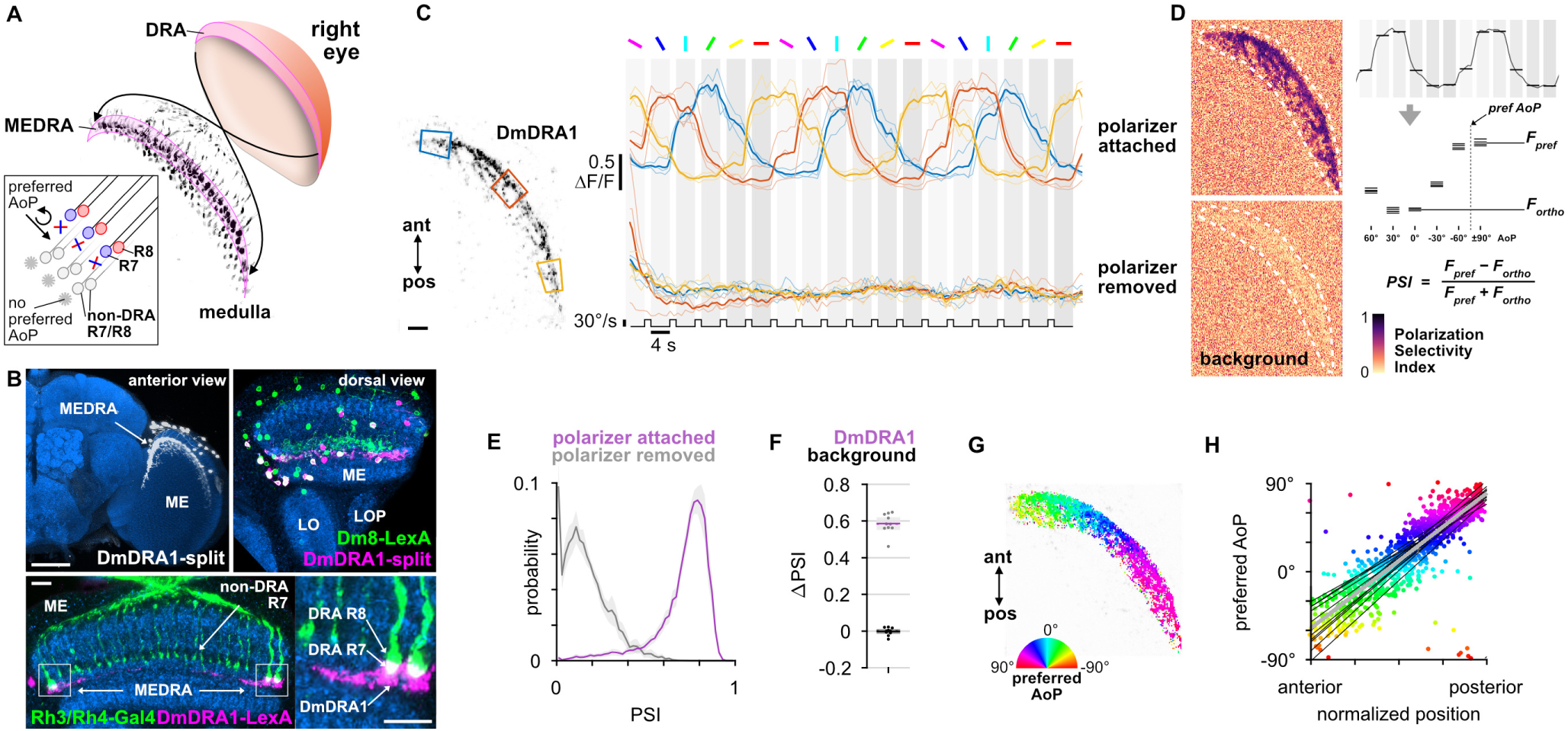
Polarized light processing in the medulla dorsal rim area. **A**: Schematic of the dorsal rim area (DRA) of the right eye and the projection of DRA R7/R8 photoreceptors to corresponding columns in the medulla dorsal rim area (MEDRA). Inset: R7 and R8 in an individual column are tuned to orthogonal angles of polarization (AoP), and their tunings change linearly across the MEDRA. **B**: Top, left: Confocal projection (anterior view) of DmDRA1 expression pattern in the MEDRA (DmDRA1-split>GFP). Scale bar denotes 50 μm. Top, right: Dual-labeling of Dm8 and DmDRA1 neurons (dorsal view) (R24F06-LexA>GFP, green; DmDRA1-split>RFP, magenta) (mean cell bodies per brain hemisphere, DmDRA: 23.13, SEM 1.16; Dm8∩DmDRA: 21.25, SEM 0.49, N = 8 animals). Bottom, left: Dorsal view of the medulla showing DRA R7/R8 photoreceptors (Rh3/Rh4-Gal4, green) and their proximity to DmDRA1 neurons (R13E04-LexA, magenta). Scale bar denotes 10 μm. Bottom, right: Enlargement of medulla dorsal rim area (MEDRA). **C**: Left: Example time-averaged maximum-intensity projection showing dorso-posterior two-photon imaging view of GCaMP activity in DmDRA1 neurons (DmDRA1-split>sytGCaMP6s). Three ROIs were manually drawn in anterior (blue), dorsal (red), and posterior (yellow) MEDRA in each recording. Scale bar denotes 10 μm. Right: Time-series of normalized mean intensity values for ROIs in equivalent positions in three animals (thin traces) and their mean (thick trace), with the polarizing filter (polarizer) attached (top) and removed (bottom). Shaded patches denote periods that the polarizer remained at a fixed orientation. **D**: Example spatial maps of polarization-selectivity index (PSI) for the example recordings in **C** with the polarizer attached (top) and removed (bottom). **E**: Probability distributions of PSI values in DmDRA1 neurons with the polarizer attached (average PSI DmDRA: 0.74, CI 0.06, N = 10 animals) and removed (average PSI DmDRA1 control: 0.16, CI 0.07, N = 7 animals). Mean ± CI. **F**: Effect of polarizer on median PSI values versus controls with polarizer removed, within DmDRA1 neurons (light dots) and background regions (dark dots) in individual animals (DmDRA, pink line: mean ΔPSI = 0.59, CI 0.06, N = 10, p < 10^−6^ t-test; background, black line: mean ΔPSI = −0.002, CI 0.02, N = 10, p = 0.82, t-test). **G**: Example polarization tuning map for DmDRA1. Preferred angles of polarization are shown for each pixel with an above-threshold PSI value using the color map shown. Pixels with a below-threshold PSI value, or falling outside an ROI drawn around the DmDRA1 population, show average intensity in grayscale. Data shown are from maximum-selectivity projections through the MEDRA. **H**: Scatter plot showing the common polarotopic organization of DmDRA1 neurons. Individual points represent pixels recorded from DmDRA1 neurons, showing their normalized horizontal position in the MEDRA and their preferred angle of polarization (AoP). Thin lines show linear-circular fits for data from individual animals with significant correlations (mean ρ = 0.89, SEM 0.06, N = 10 animals), thick line shows fit for all pooled data (ρ = 0.85, N = 10 recordings, p < 10^−6^ permutation test).

### Polarized light processing in the medulla dorsal rim area

First, we concentrated on distinct morphological forms of distal medulla (Dm) interneurons which are localized to the medulla dorsal rim area (MEDRA). Two types of these interneurons have been anatomically characterized, DmDRA1 and DmDRA2. Individual DmDRA1 neurons span approximately ten MEDRA columns and receive input exclusively from DRA R7 photoreceptors while avoiding input from non-DRA columns (Sancer et al., 2019). DmDRA2 receives exclusive input from DRA R8 photoreceptors. Due to their contact with polarization-sensitive photoreceptors, both DmDRA subtypes are thought likely to respond to polarized light (Sancer et al., 2019). To test this, we generated a split-Gal4 driver (R13E04-AD, VT059781-DBD) for a population of DmDRA neurons (Fig. 1B, top left) (Courgeon and Desplan, 2019; Jenett et al., 2012). To identify which subtype expressed this driver, we co-labeled it with an established Dm8 driver (R24F06-LexA) which is known to be expressed in DmDRA1 and not DmDRA2 (Sancer et al., 2019). We found highly overlapping expression between these drivers (Fig. 1B, top right), indicating that the split-Gal4 is predominantly expressed in DmDRA1. We confirmed that DmDRA neurons in the split-Gal4 were also in close proximity to photoreceptor terminals in the MEDRA, and found clear overlap with the proximal tip of each DRA R7/R8 pair, providing further evidence of exclusive contact with DRA R7 (Fig. 1B, bottom). Hereafter, we refer to this driver as the DmDRA1-split.

After validating a polarized light stimulus by confirming the previously characterized response properties of DRA R7/R8 (Weir et al., 2016) (Fig. S1), we recorded presynaptic calcium signals in the DmDRA1-split using GCaMP6s localized to synapses (Cohn et al., 2015) while presenting different angles of polarization (AoP) to the dorsal rim (Fig. 1C, Fig. S1). We found that the activity of DmDRA1 neurons varied with the AoP presented and followed a sinusoidal response profile typical of polarization-sensitive neurons (Heinze, 2013). To quantify the extent to which the neurons were modulated by the AoP, we calculated a polarization-selectivity index (PSI) by comparing the peak response with the response at orthogonal angles (Fig. 1D). PSI values had a minimum possible value of 0, indicating equal responses to all angles presented, and a maximum of possible value of 1, indicating maximum response to two diametrically opposite angles with zero activity at their two respective orthogonal angles. Amongst DmDRA1 neurons, we found high PSI values throughout with an average of 0.74, while background regions in each recording contained an average PSI of 0.20 (Fig. 1D,E). When we repeated the experiment with the linear polarizer removed from the stimulus device, all neurons were suppressed at the initial onset of unpolarized UV light and were no longer modulated by the rotation of the device (Fig. 1C). The PSI values of the neurons then reflected this lack of modulation, falling by approximately 80%, whereas the PSI values in the background showed no change (Fig. 1D,F).

Within the population of DmDRA1 neurons, we observed preferential responses to different angles of polarized light depending on their position in the MEDRA (Fig. 1C,G). The preferred AoP showed a linear relationship with position, which we refer to as polarotopy (Fig. 1H). Moving anterior to posterior in the right optic lobe, the preferred AoP shifted counter-clockwise (Fig. 1G,H). This polarotopy was mirrored in the left optic lobe, with a similar range of preferred AoPs represented in the opposite posterior-anterior direction (Fig. S1I). Throughout the MEDRA, the preferred AoPs of DmDRA1 neurons closely matched those of R8 photoreceptors at similar positions (Fig. 1H, Fig. S1E). Since R7/R8 are likely inhibitory (Davis et al., 2020; Gao et al., 2008), we expected that the preferred AoP of a neuron postsynaptic to either R7 or R8 would be shifted by 90°. We therefore posit that it is R7 signals that are responsible for the predominant response characteristics of DmDRA1 neurons, supporting our anatomical data (Fig. 1B) and the connectivity of the DmDRA1 subtype (Sancer et al., 2019).

We then asked whether DmDRA1 neurons are inhibited by anti-preferred angles, which would likely require antagonistic processing of local, orthogonally-tuned R7 and R8 signals in the MEDRA. Although DmDRA1 does not contact R8, inhibitory interactions between R7/R8 in each column suggest that direct input may not be necessary (Schnaitmann et al., 2018; Weir et al., 2016). We first identified anterior regions in the MEDRA where the preferred AoP of DmDRA1 was found to be around 0° in the previous tuning experiment (Fig. 1G) and generated ROIs around similarly tuned pixels (Fig. S2A,B). We then measured the responses of each ROI to flashes of UV light with 0° and 90° AoP (Fig. S2C). The preferred AoP of 0° caused an increase in activity while flashes at 90° caused inhibition of greater magnitude, followed by a slight rebound above baseline after the offset of the flash (Fig. S2C). For light flashes with the polarizer removed we observed inhibition of DmDRA1 at all regions, regardless of position in the DRA (Fig. S2C’). Taken together, these results support a model of polarization-opponent processing, whereby DmDRA1 neurons are excited and inhibited by orthogonal angles of polarized light, and inhibited by unpolarized light.

### Medulla projection neurons convey polarized light signals to the AOTU

In other insect species, polarization-sensitive photoreceptors in the dorsal rim are thought to provide input to transmedulla neurons (also referred to as line-tangential neurons) which project from the optic lobe to the anterior optic tubercle (AOTU) (Homberg et al., 2003; Immonen et al., 2017; el Jundi et al., 2011; Pfeiffer and Kinoshita, 2012; Zeller et al., 2015). In all species investigated, it is the small subunit of the AOTU (often called the lower-unit, LU) which is involved in processing polarized light signals (Heinze, 2013), although to our knowledge these signals have not been explored in transmedulla neurons themselves. In *Drosophila*, corresponding medullo-tubercular (MeTu) neurons have been described (Fig. 2A), some of which have been shown to play a role in color vision-dependent behaviors (Omoto et al., 2017; Otsuna et al., 2014). The dendrites of individual MeTu neurons typically innervate 10–15 columns of the medulla in layers M6–7 (Omoto et al., 2017) (Fig. S3) and, as an ensemble, tile larger areas of the medulla (Fig. 2A). We predicted that MeTu neurons with dendrites in the MEDRA would be postsynaptic to DmDRA1 neurons and/or DRA R7/R8, and would therefore similarly respond to polarized light.

**Figure 2:**
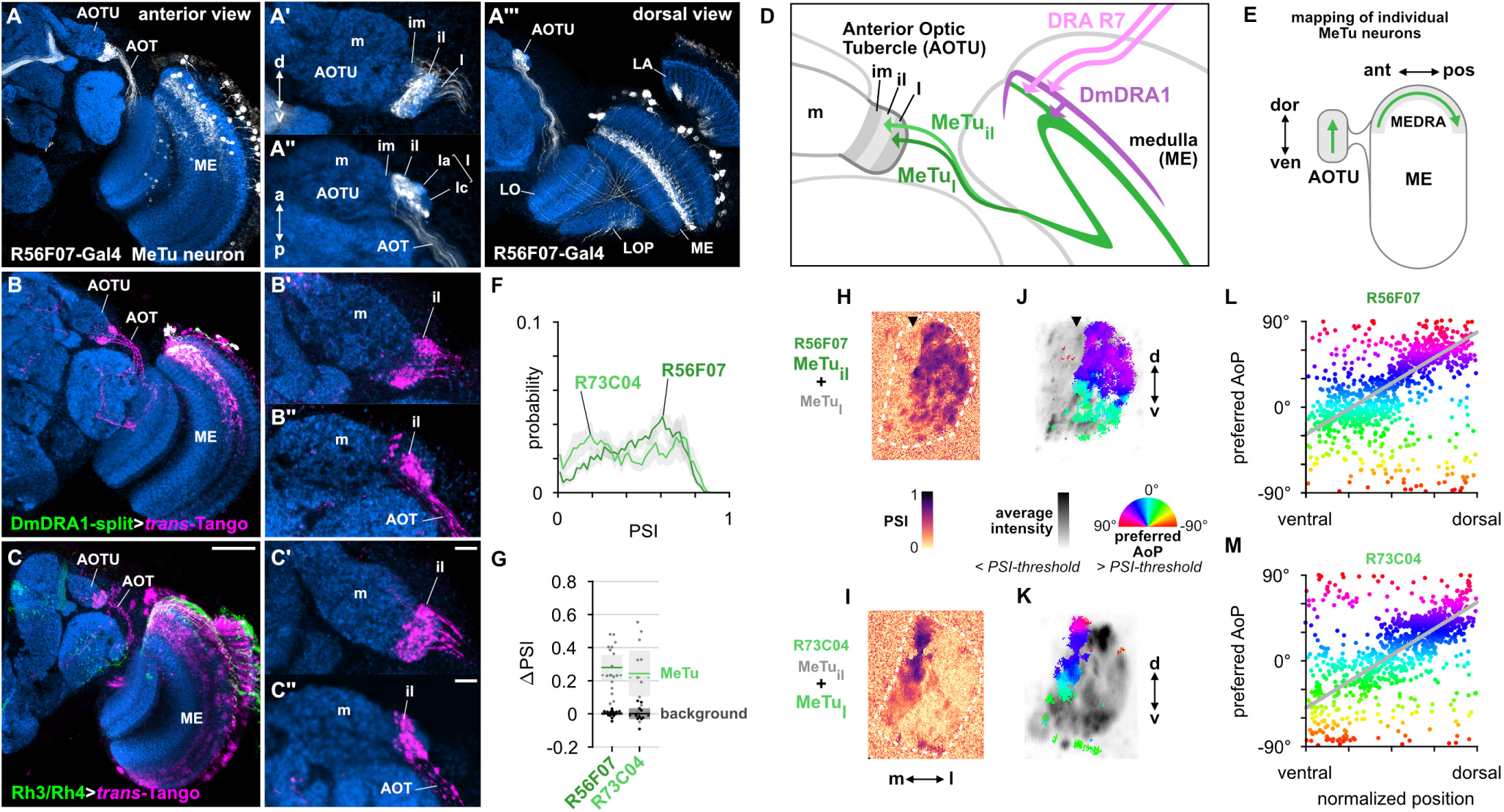
Medulla projection neurons convey polarized light signals to the AOTU. **A**: Confocal projection (anterior view) of R56F07-Gal4 driving a population of MeTu neurons with dendrites in the dorsal medulla (ME) and projections to anterior optic tubercle (AOTU) via the anterior optic tract (AOT). High magnification anterior (**A’**) and dorsal (**A’’**) views. **A’’’**: Dorsal view. **B**: Confocal projection (anterior view) of *trans*-Tango signal (magenta) labeling putative postsynaptic partners from DmDRA-Gal4 (green). High magnification anterior (**B’**) and dorsal (**B’’**) view. **C**: Confocal projection (anterior view) of *trans*-Tango signal (magenta) labeling putative postsynaptic partners from Rh3/Rh4-Gal4 (green), which labels DRA R7/R8 + non-DRA R7. Scale bar denotes 50 μm. High magnification anterior (**C’**) and dorsal (**C’’**) views (scale bars denote 10 μm). **D**: Schematic of proposed parallel connectivity in the medulla dorsal rim area (MEDRA) and regions of the AOTU targeted by polarization-sensitive MeTu neurons. **E**: Schematic of proposed one-dimensional mapping of MEDRA position to AOTU based on single-cell clones (see Fig. S3). **F**: Probability distributions of PSI values in MeTu neurons (average PSI R56F07: 0.48, CI 0.14, N = 17 animals; R73C04: 0.42, CI 0.20, N = 11 animals). Mean ± CI. **G**: Effect of polarizer on median PSI values versus controls with polarizer removed, within MeTu neurons (light dots) and background regions (dark dots) in individual animals (R56F07 MeTu, green line: mean ΔPSI = 0.28, CI 0.14, N = 17, p < 10^−6^ t-test; R56F07 background, black line: mean ΔPSI = 0.001, CI 0.02, N = 17, p = 0.84, t-test; R73C04 MeTu, green line: mean ΔPSI = 0.25, CI 0.20, N = 11, p = 0.03 t-test; R73C04 background, black line: mean ΔPSI = 0.001, CI 0.05, N = 11, p = 0.96, t-test). **H**: Example spatial map of polarization-selectivity index (PSI) in MeTu terminals in the AOTU (R56F07-Gal4>sytGCaMP6s; predominantly MeTu_il_ neurons innervating intermediate-lateral (il) domain, with smaller proportion of MeTu_la_ innervating lateral-anterior (la) domain, see **A’**’). Arrowhead indicates medial region of population with low PSI values cf. average activity in **J**. **I**: Example spatial map of PSI in MeTu terminals in the AOTU for an alternative driver (R73C04-Gal4>sytGCaMP6s; predominantly MeTu_l_ neurons innervating lateral (l) domains, with smaller proportion of MeTu_il_ innervating intermediate-lateral (il) domain, see Fig. 3G’). **J**: Example polarization tuning map for above-threshold pixels in R56F07 MeTu neurons from the example recording in **H. K**: As in **J**, for R73C04 MeTu neurons from the example recording in **I**. **L**: Scatter plot showing the predominant polarotopic organization of R56F07 MeTu neurons. Individual points represent pixels recorded in MeTu neurons, showing their normalized vertical position in the MEDRA and their preferred angle of polarization (AoP). Line shows fit for all pooled data (ρ = 0.68, N = 7 animals, p < 10^−6^ permutation test). **M**: As in **L**, for R73C04 MeTu neurons (ρ = 0.58, N = 10 animals, p < 10^−6^ permutation test).

We used the anterograde circuit tracing technique *trans*-Tango (Talay et al., 2017) to identify putative postsynaptic partners of the DmDRA1 neurons and R7/R8 photoreceptors (Fig. 2B,C). We found that DmDRA1-split driving *trans*-Tango labeled a population of neurons in the dorsal medulla, along with innervation of the small, lateral subunit of the AOTU via a fiber bundle in the anterior optic tract (AOT) (Fig. 2B), which matched the anatomy of MeTu neurons (Fig. 2A). We then used a Gal4 driver which targets neurons expressing the UV-sensitive rhodopsins Rh3 and Rh4 (pan-R7-Gal4, which we refer to as Rh3/Rh4-Gal4), which includes DRA R7/R8, and again found *trans*-Tango labeling of the small subunit of the AOTU (Fig. 2C). However, since the Rh3/Rh4 driver is also expressed in non-DRA R7 photoreceptors (Fig. 2C), the labeling of MeTu neurons we observed could have been due to synaptic contacts exclusively outside of the MEDRA. To evaluate this possibility, we co-labeled a population of MeTu neurons and all photoreceptors using the antibody mAb24B10 (Fujita et al., 1982) (Fig. S3A). Throughout layer M6 in the dorsal medulla, MeTu dendrites were in close proximity to R7/R8 terminals and we found clear overlap with R7 terminals in the MEDRA (Fig. S3A). In short, these putative connections suggest a parallel pathway for polarization signals in the MEDRA: DRA R7→DmDRA1, DmDRA1→MeTu, DRA R7→MeTu.

Several discrete populations of MeTu neurons have been characterized, based on the distinct domains of the small subunit of the AOTU that their terminals occupy: intermediate-medial (im), intermediate-lateral (il), and lateral (l), which is further divided into anterior (la), central (lc), and posterior (lp) domains (Fig. 2A’,A’’, Fig. S3B). The larger subunit comprising the medial domain (m) is not innervated by MeTu neurons and corresponds to the polarization-insensitive upper-unit (UU) of other species (Omoto et al., 2017; Timaeus et al., 2017). We examined the domains of the AOTU targeted by the putatively polarization-sensitive MeTu neurons which were labeled by *trans*-Tango (Fig. 2B’–C’). Both the DmDRA1 and Rh3/Rh4 *trans*-Tango experiments predominantly labeled the intermediate-lateral domain (AOTU_il_), with encroachment on the lateral domain (AOTU_l_) (Fig. 2B’’–C’’). We found no detectable intermediate-medial (AOTU_im_) or medial (AOTU_m_) labeling in either (Fig. 2B’–C’). We next identified two Gal4 drivers for populations of MeTu neurons arborizing in the AOTU_l_ and AOTU_il_: one with dendrites predominantly tiling the dorsal medulla (R56F07-Gal4) (Fig. 2A) and one with dendrites throughout the medulla (R73C04-Gal4) (Fig. 3G) (Omoto et al., 2017). From confocal images of single-cell MCFO (MultiColor FlpOut) clones (Nern et al., 2015), we determined a consistent relationship between the anterior→posterior position of MeTu dendrites in the MEDRA and the ventral→dorsal position of MeTu axon terminals in the AOTU (Fig. 2E, Fig. S3). For MeTu neurons with dendrites outside of the MEDRA, we found no clear relationship between ventrodorsal position in the medulla and mediolateral position in the AOTU, confirming a previous study (Timaeus et al., 2017).

**Figure 3:**
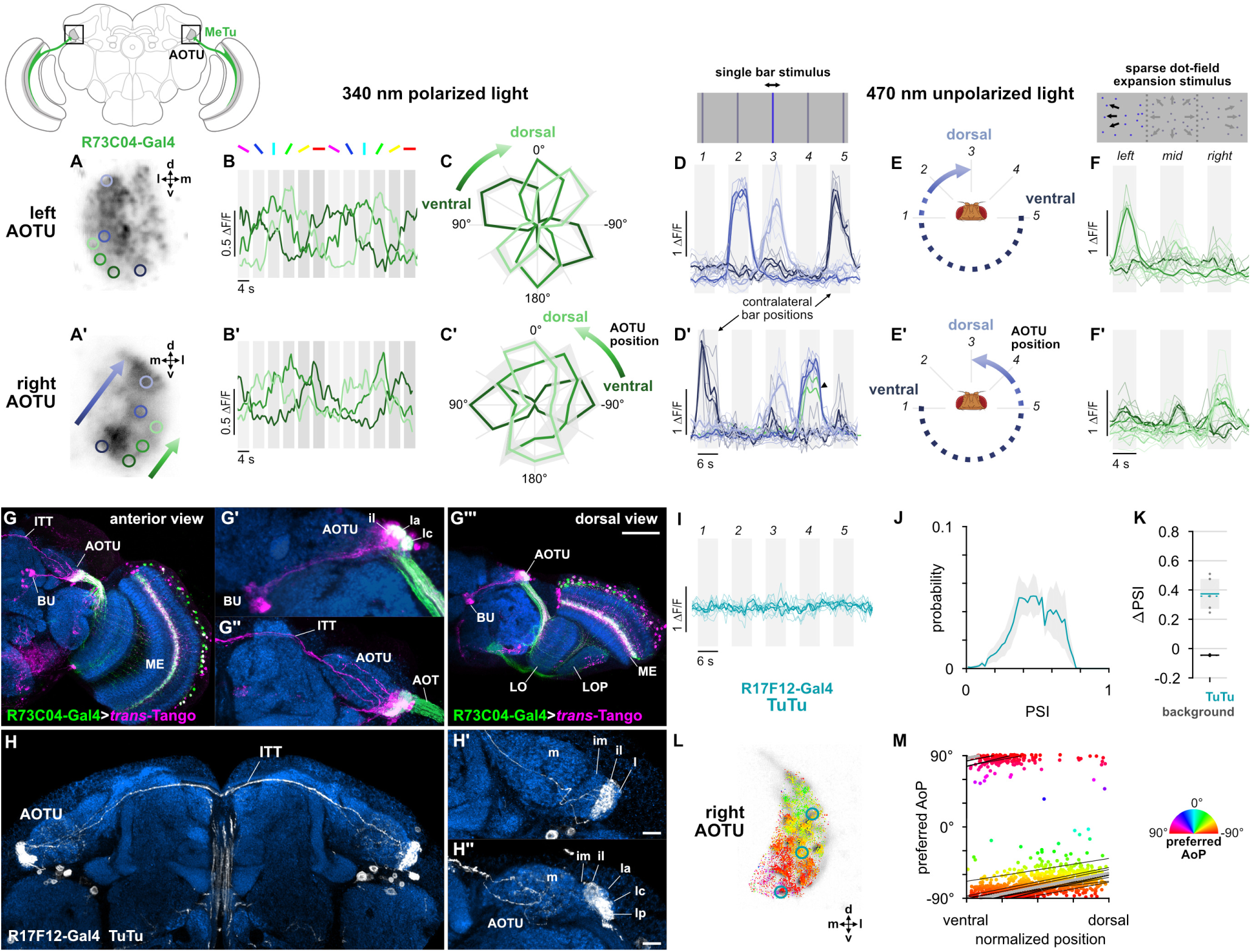
Visual features encoded in the AOTU and bilateral interactions. **A**: Example time-averaged maximum-intensity projection showing GCaMP activity in R73C04 MeTu neurons in the AOTU and examples of lateral ROIs (green) and medial ROIs (blue) (R73C04-Gal4>sytGCaMP6s). **B**: GCaMP activity in lateral MeTu neurons showing responses to different angles of polarization. Each trace shows the mean of ROIs at equivalent positions in three different animals (one ROI per animal). **C**: Normalized tuning curves for responses shown in B. Mean ± SEM. **D**: Responses of MeTu neurons in medial positions to an unpolarized blue bar oscillating in five positions in the frontal visual field. Traces of the same color are from ROIs in equivalent positions in the AOTU in three different animals, thick traces show their mean. Bar positions 1 and 5 correspond to ± 90° azimuth in the contralateral visual field for recordings in the right (**D**’) and left (**D**) AOTU, respectively. Arrowhead in D’ indicates the response of an ROI in a lateral position (green) with similar responses to the bar stimulus. **E**: Proposed mapping of azimuthal position in visual field to vertical position in AOTU, based on **D**. **F**: Responses of MeTu neurons in lateral positions to a sparse dot-field expansion pattern presented in three regions of the frontal visual field. Traces of the same color are from ROIs in equivalent positions in the AOTU in three animals, thick traces show their mean. **G**: Confocal projection (anterior view) of *trans*-Tango signal (magenta) labeling putative postsynaptic partners of R73C04-Gal4 MeTu neurons (green). **G’**: High magnification dorsal view highlighting TuBu neurons projecting from AOTU to bulb (BU). **G’’**: High magnification anterior view highlighting projections to contralateral AOTU. **G’’’**: Dorsal view. Scale bar denotes 50 μm. **H**: Confocal projection (anterior view) of TuTu neuron expression pattern (R17F12-Gal4>GFP). High magnification anterior (**H’**) and dorsal (**H’’**) views. Scale bars denote 10 μm. **I**: As in **D**, for TuTu neurons. **J**: Probability distribution of PSI values in TuTu neurons (average PSI TuTu: 0.48, CI 0.12, N = 5 animals). Mean ± CI. **K**: Effect of polarizer on median PSI values versus controls with polarizer removed, within TuTu neurons (light dots) and background regions (dark dots) in individual animals (TuTu, blue line: mean ΔPSI = 0.34, CI 0.12, N = 5, p = 0.02 t-test; background, black line: mean ΔPSI = −0.045, CI 0.05, N = 5, p < 10^−4^ t-test). **L**: Example polarization tuning map for above-threshold pixels in the terminals of R17F12 TuTu neurons in a single imaging plane (R17F12-Gal4>sytGCaMP6s). **M**: Scatter plot showing the predominant polarotopic organization of R17F12 TuTu neurons. Thin lines show linear-circular fits for data from individual animals with significant correlations (mean ρ = 0.65, SEM 0.06, N = 5 animals), thick line shows fit for all pooled data (ρ = 0.56, N = 5 recordings, p < 10^−6^ permutation test).

We recorded presynaptic calcium signals in the AOTU for the two MeTu drivers in response to rotations of the polarizer, as in Fig. 1. In both MeTu populations, we found broader PSI distributions (Fig. 2F) than in the DmDRA1 neurons recorded in the MEDRA (Fig. 1E). Nonetheless, compared to control experiments with the polarizer removed, the polarizer caused a statistically significant increase in average PSI values in both MeTu distributions (Fig. 2G). We observed that the highest PSI values were spatially restricted to a vertical band within the AOTU (Fig. 2H,I), indicating that MeTu terminals which were strongly modulated by the polarization stimulus occupied a common region, while adjacent regions contained terminals which were generally modulated less. We surmise that these regions of differing polarization-sensitivity result from each population containing a combination of MeTu neurons with dendrites contacting the MEDRA, which constitutes only around 5% of medulla columns (Weir et al., 2016), and neurons with dendrites outside the MEDRA. We also note the proportion of PSI values below 0.5 was slightly lower in the population containing neurons with dendrites in the dorsal medulla only (R56F07) compared to the ventral and dorsal population (R73C04) (Fig. 2F,H,I). In R56F07, the most responsive MeTu terminals were found within the most lateral regions of the population in the AOTU (Fig. 2H, Fig. S3E). In R73C04, the most responsive terminals tended to be clustered in a narrow medial band of the population (Fig. 2I, Fig. S3F), likely corresponding to the anterior region of AOTU_il_ and possibly AOTU_la_.

Based on the polarotopic organization of R7/R8 and DmDRA1 in the MEDRA, as well as the mapping of MEDRA to AOTU by MeTu neurons (Fig. 2E), we predicted that polarization-sensitive MeTu neurons would exhibit a counter-clockwise shift in their preferred AoP from ventral to dorsal in the right AOTU. To assess this, we examined pixels with above-threshold PSI values (>1 SD greater than the mean background value, see Methods), which limited the analysis to polarization-sensitive MeTu terminals (Fig. 2J,K). Across animals, both populations showed a predominant polarotopic organization which matched our prediction: from ventral to dorsal in the right AOTU, the preferred AoP shifted counter-clockwise (Fig. 2L,M). This polarotopy is consistent with MeTu neurons receiving polarized light responses from either DmDRA1 or DRA R7 in the MEDRA and conveying them to the AOTU with the positional mapping we identified (Fig. 2D,E). Consistent with this mapping, we observed no clear relationship between preferred AoP and horizontal position (Fig. S3E,F). However, we observed vertical organizations of responses which deviated from the norm in approximately 20% of recordings across both drivers. The most common of these resembled an inverted form of the predominant polarotopy (from ventral to dorsal in the right AOTU, the preferred AoP rotated clockwise) and also typically contained tunings to a different range of AoPs than the predominant organization (Fig. S3I,I’). Although we could not determine why one organization was observed over another, this finding suggests that a further transformation of MeTu responses may take place. However, a reversed mapping of responses could be achieved by combining signals originating from the contralateral eye (Fig. S1G,H), which we explore below.

### Visual features encoded in the AOTU and bilateral interactions

We wondered whether functional divisions of MeTu responses exist within the AOTU, which might contain different polarotopic organizations or spatially segregated responses to unpolarized visual features not mediated by the MEDRA. We first examined the spatial organization of polarized light responses in regions which contained low or below-threshold PSI values in the previous experiment (Fig. 2I,K). Within lateral MeTu terminals in R73C04 likely occupying the ventral AOTU_lc_ domain (green ROIs, Fig. 3A), we found moderate modulation of activity during the rotation of the polarizer (Fig. 3B). Similar to the terminals with above-threshold PSI values (Fig. 2K), we observed a vertical polarotopic organization consistent with the anatomical mapping of MeTu neurons (Fig. S3B–D): in a dorsal direction, the AoP rotated counter-clockwise in the right AOTU and clockwise in the left AOTU (Fig. 3C). We then recorded MeTu responses to unpolarized, small-field vertical bar stimuli at different positions in the visual field (Fig. 3D). Within an intermediate band of MeTu terminals likely corresponding to AOTU_la_ (blue ROIs, Fig. 3A), we observed clear responses to bars in ipsilateral-frontal and frontal positions, with the more frontal position represented dorsally in the AOTU on both sides of the brain (Fig. 3D). In the ventral AOTU, we found responses to bars presented in the contralateral-lateral visual field (± 90° azimuth), outside the field of view of the ipsilateral eye (Fig. 3D,E). Together, these results suggest that the AOTU contains retinotopic representations of visual space and angles of polarization within different regions (Fig. 3C,E). Furthermore, these regions do not appear to be mutually exclusive, as we occasionally observed responses to both polarized and unpolarized stimuli at the same location (green trace, Fig. 3D’). For example, MeTu terminals in regions which were modulated by the polarizer (green ROIs, Fig. 3A) also responded to a wide-field optic-flow pattern presented at different locations (Fig. 3F), further highlighting the range of visual features represented in a particular region of the AOTU.

Evidence from other insects suggested that we might find bilateral, inter-tubercle neurons which, if in contact with MeTu neurons, could be conveying the responses we observed in the AOTU to contralateral stimuli (Heinze et al., 2013; Pfeiffer and Kinoshita, 2012; Pfeiffer et al., 2005). We used the MeTu driver R73C04-Gal4 to drive *trans*-Tango and reveal putative postsynaptic neurons in the AOTU (Fig. 3G). We found clear labeling of a population of neurons projecting to the bulb which resembled the tubercular-bulbar (TuBu) neurons (Omoto et al., 2017) (Fig. 3G’), in addition to labeling of the inter-tubercle tract (ITT) (Strausfeld, 1976) (Fig. 3G’’), suggesting inter-hemispheric signalling postsynaptic to MeTu neurons in the AOTU. We then identified a Gal4 driver (R17F12-Gal4) that is expressed in a population of two tubercular-tubercle (TuTu) neurons per brain hemisphere, with axonal projections to the contralateral AOTU via the ITT (Fig. 3H). Within the AOTU, these TuTu neurons predominantly innervate the intermediate-lateral domain (AOTU_il_) (Fig. 3H’). We recorded presynaptic calcium activity in the terminals of contralateral TuTu neurons in the AOTU (Fig. 3I,J). Unexpectedly, we did not find responses to the unpolarized bar stimuli at any of the positions tested (Fig. 3I), indicating that these TuTu neurons likely do not mediate the contralateral responses we observed in the MeTu neurons (Fig. 3D). Rather, we found that the TuTu neurons were polarization-sensitive with PSI values similar to those of the MeTu neurons (Fig. 3K,L), and tunings to a limited range of polarization angles (∼30°) centered around a near-horizontal orientation (Fig. 3L,M). Therefore, the anatomy, polarization-sensitivity, and number of TuTu neurons suggests that they may correspond to the TuTu1 neurons described in locusts, although their preferred AoPs differ (Pfeiffer et al., 2005). TuTu1 neurons in the locust have also been shown to respond to unpolarized visual stimuli, however their responses were also selective for both spatial position and color, and the unpolarized stimuli presented here are not directly comparable (Pfeiffer and Homberg, 2007). The specificity of TuTu1 responses is thought to reflect their role in time-compensated processing of polarized light signals and the integration of information about the position of the sun and spectral content of the sky.

### A population of TuBu neurons receives polarized light signals in the AOTU

Next, we focused on the TuBu neurons and asked whether they receive polarization signals in the lateral (l) and intermediate-lateral (il) domains of the anterior optic tubercle (AOTU), as suggested by *trans*-Tango labeling from polarization-sensitive MeTu neurons (Fig. 3G). We examined three populations of TuBu neurons, grouped according to the region of the bulb (BU) they project to: superior (TuBu_s_), inferior (TuBu_i_), and anterior (TuBu_a_) (Fig. 4A). The dendrites of TuBu neurons in each population have also been shown to predominantly innervate stereotypical domains of the AOTU (Omoto et al., 2017) (Fig. 4A). We recorded calcium activity using Gal4 drivers for each population, noting that the driver for superior bulb-projecting TuBu_s_ neurons (R88A06-Gal4) is also expressed in TuBu_a_ neurons. Among the dendrites of TuBu neurons recorded in the AOTU, we found that the populations innervating the AOTU_l_ and AOTU_il_ domains (TuBu_s_ and TuBu_a_, respectively) contained high PSI values that indicated strong modulation by the polarizer (Fig. 4B), with average values significantly higher than the background regions of recordings (Fig. 4C). In contrast, dendrites innervating the AOTU_im_ domain (TuBu_i_) contained PSI values not greater than 0.5 (Fig. 4B) and, on average, were indistinguishable from background regions (Fig. 4C). We typically found very few pixels with above-threshold PSI values in recordings of TuBu_i_ dendrites (Fig. 4D) and across all recordings we did not find a common relationship between the preferred angle of polarization (AoP) of TuBu_i_ neurons and their ventral-dorsal position within AOTU_im_ (Fig. 4E).

**Figure 4:**
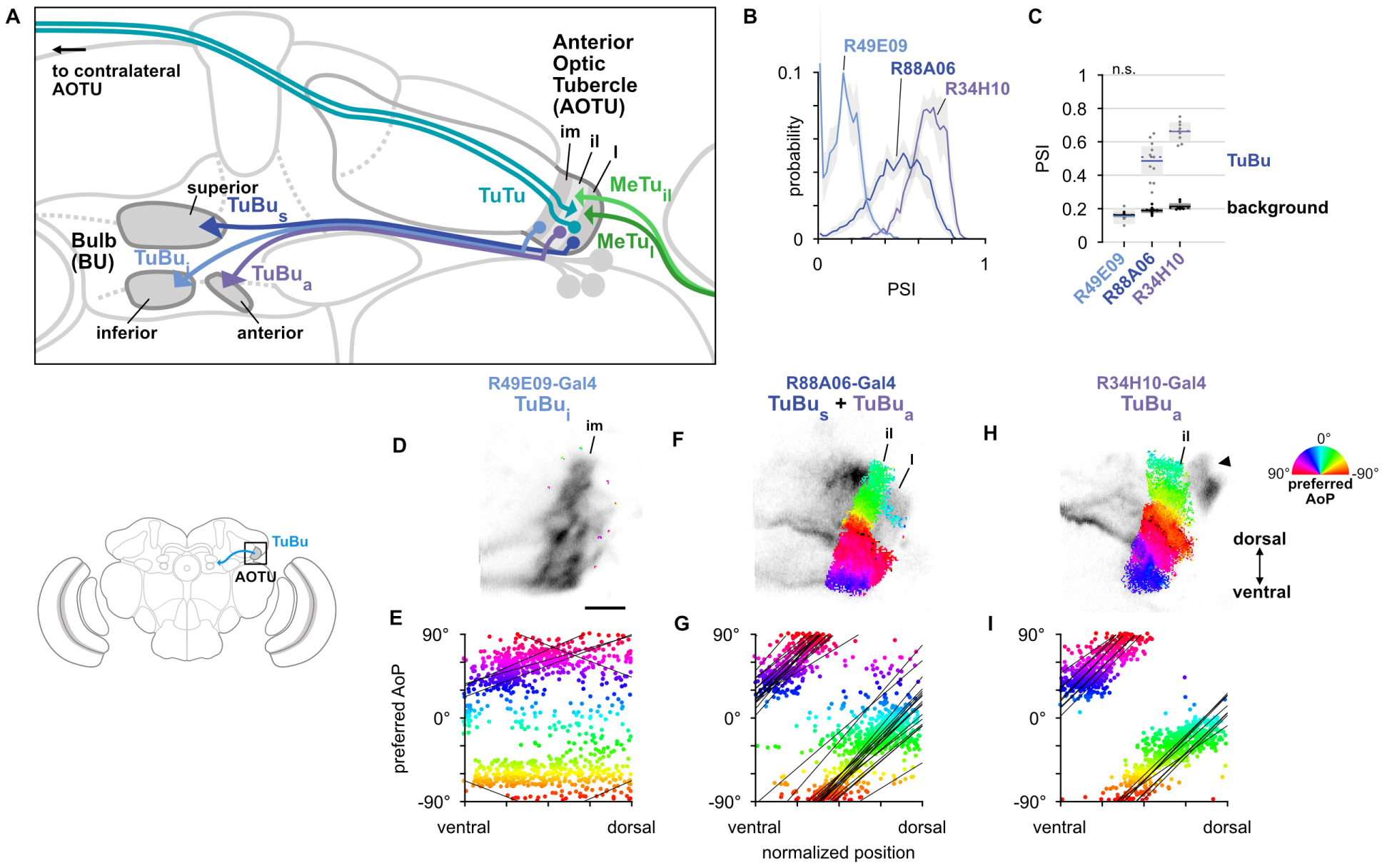
A population of TuBu neurons receives polarized light signals in the AOTU. **A**: Schematic of TuBu neuron types projecting to the bulb (BU) and connectivity in the AOTU. **B**: Probability distribution of PSI values in TuBu neurons recorded in the AOTU. Mean ± CI. Summarized in **C**. **C**: Average PSI values within TuBu neurons (light dots) and background regions (dark dots) in individual animals (**TuBu**_**i**_ neurons: 0.15, CI 0.04, background: 0.16, CI 0.14, N = 5 animals, p = 0.76 t-test; **TuBu**_**s**_ **+ TuBu**_**a**_ neurons: 0.49, CI 0.12, background: 0.19, CI 0.02, N = 11 animals, p < 10^−4^ t-test; **TuBu**_**a**_ neurons: 0.67, CI 0.06, background: 0.21, CI 0.02, N = 5 animals, p < 10^−6^ t-test). Shaded box denotes Bonferroni corrected 95% confidence interval. **D**: Example polarization tuning map for above-threshold pixels in the dendrites of TuBu_i_ neurons in a single imaging plane (R49E09-Gal4>GCaMP6s). Below-threshold pixels display average intensity in grayscale. Scale bar denotes 5 μm. **E**: Scatter plot showing the lack of polarotopic organization in TuBu_i_ neurons. Individual points represent pixels recorded from TuBu neurons, showing their normalized vertical position in the AOTU and their preferred angle of polarization (AoP). Thin lines show linear-circular fits for data from individual animals with significant correlations (mean individual ρ = 0.28, SEM 0.29, N = 4 animals; pooled data ρ = 0.19, N = 5 recordings, p < 10^−6^ permutation test). **F**: As in **D**, for a population containing TuBu_s_ and TuBu_a_ neurons (R88A06-Gal4>GCaMP6s). **G**: As in **E**, for the common polarotopic organization in TuBu_s_ and TuBu_a_ neurons (mean individual ρ = 0.63, SEM 0.21, N = 11 animals; pooled data ρ = 0.09, N = 11 recordings, p < 10^−6^ permutation test). **H**: As in **D**, for TuBu_a_ neurons (R34H10-Gal4>GCaMP6s). Arrowhead indicates cell bodies excluded from analysis. **I**: As in **E**, for the common polarotopic organization in TuBu_a_ neurons (mean individual ρ = 0.51, SEM 0.32, N = 8 animals; pooled data ρ = 0.64, N = 8 recordings, p < 10^−6^ permutation test).

Within the joint population of TuBu_s_ and TuBu_a_ neurons (R88A06-Gal4), the lateral domain (AOTU_l_) containing TuBu_s_ dendrites typically exhibited a mixture of below-threshold PSI values and a smaller proportion of above-threshold values (Fig. 4F), whereas the more-medial AOTU_il_ domain containing TuBu_a_ dendrites consistently showed above-threshold PSI values (Fig. 4F). Pooling data from both domains, the preferred AoP covered a range of angles from −90° to +90° and we found a common relationship between preferred AoP and ventral-dorsal position within the AOTU (Fig. 4G). Correspondingly, dendritic regions specifically within the population of TuBu_a_ neurons (R34H10-Gal4) contained entirely above-threshold PSI values (Fig. 4H) and obeyed the same polarotopic organization (Fig. 4I).

For the dendrites of TuBu_a_ and TuBu_s_ neurons, we found that the direction of polarotopy in the AOTU (a counter-clockwise rotation of preferred AoP from ventral to dorsal) matched the polarotopy in the putatively presynaptic MeTu neurons. However, the relative positions of tunings along the ventrodorsal axis of the AOTU do not correspond directly. For example, in the dorsal half of the AOTU the preferred AoPs of MeTu terminals were in the range 0° to +90° (Fig. 2L,M), whereas for TuBu_a_ dendrites in the dorsal half of the AOTU preferred AoPs were in the range −90° to 0° (Fig. 4I). If MeTu neurons are indeed presynaptic to TuBu neurons in the AOTU, this result suggests either inhibitory input from MeTu neurons, which would effectively shift the preferred AoP by 90°, or the integration of additional inputs from unidentified polarization-sensitive elements at TuBu dendrites.

### The anterior bulb is an entry point for polarized light signals into the central complex

We next asked how responses of TuBu neurons are organized in the bulb (BU). As in other insects, the BU features giant synapses (‘micro-glomeruli’) formed by TuBu endings and their targets, the ring neurons. In *Drosophila*, the BU consists of three anatomical regions: superior (BUs), inferior (BUi), and anterior (BUa) (Fig. 4A). We recorded presynaptic calcium activity in the micro-glomerular terminals of TuBu neuron populations that target each region. We first examined the prevalence of polarization-modulated activity, indicated by the polarization-selectivity index (PSI). Spatial maps of PSI values revealed that the majority of TuBu_s_ neurons recorded in micro-glomeruli in the BUs contained low PSI values, and interspersed among them were micro-glomeruli with high PSI values (Fig. 5A). The mixture of polarization-sensitive and insensitive micro-glomeruli is conveyed by the broad distribution, skewed towards zero, of PSI values found across all pixels recorded in the BUs (Fig. 5B). In contrast, the narrow distribution of PSI values close to zero in BUi micro-glomeruli demonstrates the absence of polarization-sensitive TuBu_i_ neurons (Fig. 5B). Finally, we found that all TuBu_a_ neurons recorded exhibited high PSI values in the BUa (Fig. 5A,B), in two Gal4 drivers. Average PSI values in the BUa were greater than 0.5 in both drivers (Fig. 5C), while in the BUi and BUs, the average PSI values were not significantly different from the average in background regions of recordings, typically around 0.2 (Fig. 5C).

**Figure 5:**
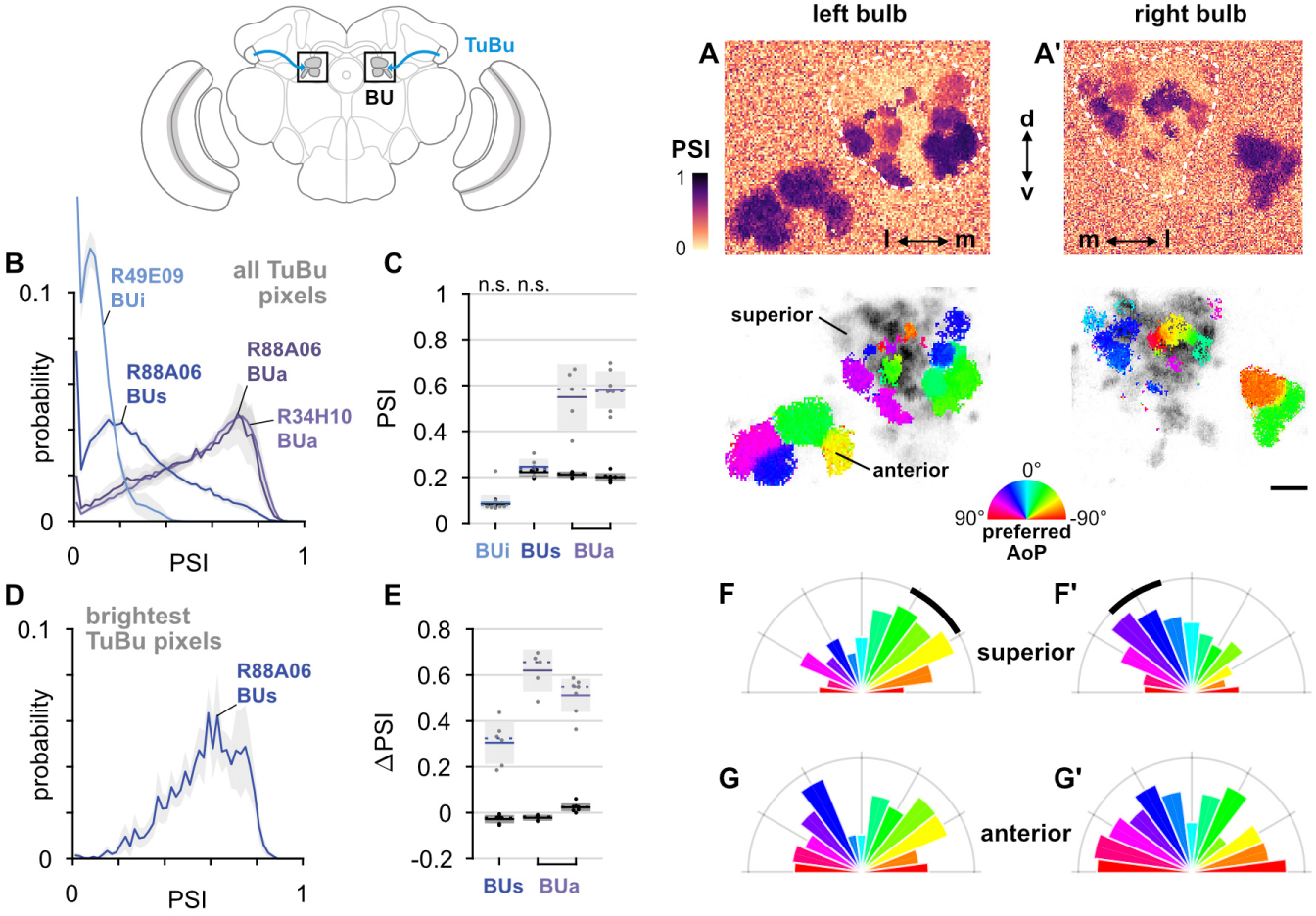
The anterior bulb is an entry point for polarized light signals into the central complex. **A**: Example spatial maps of polarization-selectivity index (PSI, top) and tuning (bottom) in TuBu neuron output micro-glomeruli in the superior and anterior regions of the left (**A**) and right (**A’**) bulbs in an individual fly (R88A06-Gal4>sytGCaMP6s). Scale bar denotes 5 μm. **B**: Probability distribution of PSI values in all pixels recorded in TuBu neurons in the three regions of the bulb (BU). Mean ± CI. Summarized in **C**. **C**: Average PSI values within TuBu neurons in the BU (light dots) and background regions (dark dots) in individual animals (**BUi**: 0.09, CI 0.04, background: 0.09, CI 0.08, N = 12 animals, p = 0.68 t-test; **BUs**: 0.25, CI 0.04, background: 0.21, CI 0.02, N = 6 animals, p = 0.18 t-test; **BUa** (R88A06): 0.59, CI 0.10, background: 0.21, CI 0.01, N = 5 animals, p = 0.0002 t-test; **BUa** (R34H10): 0.58, CI 0.09, background: 0.20, CI 0.02, N = 7 animals, p < 10^−4^ t-test). Shaded box denotes Bonferroni corrected 95% confidence interval. **D**: Probability distribution of PSI values in 10% brightest pixels recorded in TuBu_s_ neurons in BUs. Mean ± CI. Summarized in **E**. **E**: Effect of polarizer on median PSI values versus controls with polarizer removed, within TuBu neurons (light dots) and background regions (dark dots) in individual animals (mean ΔPSI **TuBu**_**s**_ neurons: 0.31, CI 0.09, N = 6, p = 0.0005 t-test, background: −0.03, CI 0.02, N = 6, p = 0.02, t-test; **TuBu**_**a**_ neurons (R88A06): 0.62, CI 0.09, N = 5, p < 10^−4^ t-test, background: −0.022, CI 0.09, N = 5, p = 0.18, t-test; **TuBu**_**a**_ neurons (R34H10): 0.51, CI 0.08, N = 7, p < 10^−5^ t-test, background: −0.023, CI 0.02, N = 7, p = 0.19, t-test). Shaded box denotes Bonferroni corrected 95% confidence interval. **F**: Polar histogram of preferred angles of polarization in TuBu_s_ neurons recorded in the left (**F**) and right (**F’**) superior bulb. Normalized probabilities in each bin are displayed as area of wedge; radial lengths of wedges not directly comparable. Arc denotes mean resultant angle ± 95% confidence interval (**TuBu**_**s**_ left: 0.36 −42.4° CI 16.6°, N = 4, p = 0.002 Rayleigh uniformity test; **TuBu**_**s**_ right: 0.31 30.3° CI 15.1°, N = 5, p = 0.0006 Rayleigh uniformity test). **G**: As in **F**, for TuBu_a_ neurons recorded in the anterior bulb (R34H10) (**TuBu**_**a**_ left: 0.08 −60.6° CI N/A, N = 6, p = 0.62 Rayleigh uniformity test; **TuBu**_**a**_ right: 0.14 −66.0° CI N/A, N = 6, p = 0.22 Rayleigh uniformity test).

We further explored the PSI values in the BUs by isolating the brightest pixels in TuBu_s_ neurons in each recording, which were likely to represent active neurons (Fig. 5D). We found that the distribution of PSI values among the brightest pixels was shifted towards one and was qualitatively different to the distribution across all pixels (Fig. 5B,D). We then compared the average PSI value of the brightest pixels in the BUs with their average value in control experiments with the polarizer removed, and repeated this procedure with the brightest pixels in the BUa as a reference. Among active pixels in both the BUs and BUa we found a significant effect of the polarizer on PSI values versus controls, with the effect size larger in the latter (Fig. 5E). In sum, we found polarized light responses in TuBu neuron output micro-glomeruli in both the superior and anterior bulb, and no appreciable responses to polarized light in TuBu neuron outputs in the inferior bulb. We interpret these findings as being consistent with the corresponding dendritic responses of TuBu neurons in the AOTU (Fig. 4B).

We then asked whether the information about polarized light available in the BUs and BUa differed in some way, for example by encoding different ranges of angles. We observed that a cluster of micro-glomeruli towards the medial edge of the superior bulb tended to show preferential responses to similar angles of polarization (AoP) (Fig. 5A, bottom). When we examined the distribution of preferred AoPs in the BUs we found a non-uniform distribution with the highest frequency of preferred AoPs around −45° in the left bulb (Fig. 5F) and +45° in the right bulb (Fig. 5F’). In the anterior bulb (BUa) on both sides, we found an approximately uniform representation of preferred AoPs in TuBu_a_ neurons (Fig. 5G, G’). We expected that a uniform representation of the full range of polarization space would be necessary for decoding heading direction from skylight polarization patterns. The over-representation of certain AoPs in BUs micro-glomeruli resembles a detector for a particular feature, such as horizontally polarized reflections from the surface of water, rather than the main input to a system for polarized light-based navigation. Upon inspection, we did not see a clear linear organization of preferred AoPs in either the BUs or the BUa, a marked contrast to the consistent organization in TuBu dendrites in the AOTU (Fig. 4H,I). Circular organizations of TuBu neurons in the bulb have been proposed (Timaeus et al., 2017) and we explore these in the BUa in the next section (Fig. S5).

TuBu neurons have previously been shown to respond to unpolarized visual stimuli presented to regions of the eye outside the DRA (Omoto et al., 2017; Shiozaki and Kazama, 2017; Sun et al., 2017). To compare the responses of the three groups of TuBu neurons, we presented a wide-field flash of unpolarized blue light and recorded responses in each population in the AOTU and BU (Fig. S4A). TuBu_s_ and TuBu_i_ neuron populations showed responses to the flash in the AOTU and, more strongly, in the BU, while TuBu_a_ neurons recorded in either neuropil were inhibited by the unpolarized light stimulus (Fig. S4). We note that prior work appeared to show excitation of BUa micro-glomeruli in response to unpolarized small-field stimuli presented in the contralateral visual field and inhibition in response to ipsilateral stimuli (Shiozaki and Kazama, 2017). These results may reflect excitatory and inhibitory receptive fields of TuBu_s_ neurons, while our recordings indicate that inhibition dominates the response of the population to wide-field visual stimuli.

### R4m ring neurons receive polarization-tuned responses from TuBu neurons

Taken together, our recordings of TuBu neurons indicate that polarized light signals are potentially delivered to the central complex via two parallel pathways: one through the superior bulb (BUs), containing a limited representation of polarization space in addition to other visual information, and a second channel through the anterior bulb (BUa). In the bulb, TuBu neuron presynaptic terminals innervate the globular dendrites of ring neurons in a largely one-to-one fashion, forming individual micro-glomeruli. Ring neurons project medially to the ellipsoid body (EB) (Fig. 6A), where their arborizations have a circular form and are both dendritic and axonal (Fig. 6B) (Hanesch et al., 1989; Omoto et al., 2018). We recorded calcium activity in the dendrites of two populations of ring neurons in the bulb, one innervating the medial two-thirds of the BUs (R2; R19C08-Gal4) and one innervating the BUa (R4m; R34H10-Gal4) (Fig. 6A). Both R2 and R4m ring neuron populations target the outer central domain of the EB, albeit following different trajectories (Fig. 6A,B) (Omoto et al., 2017, 2018).

**Figure 6:**
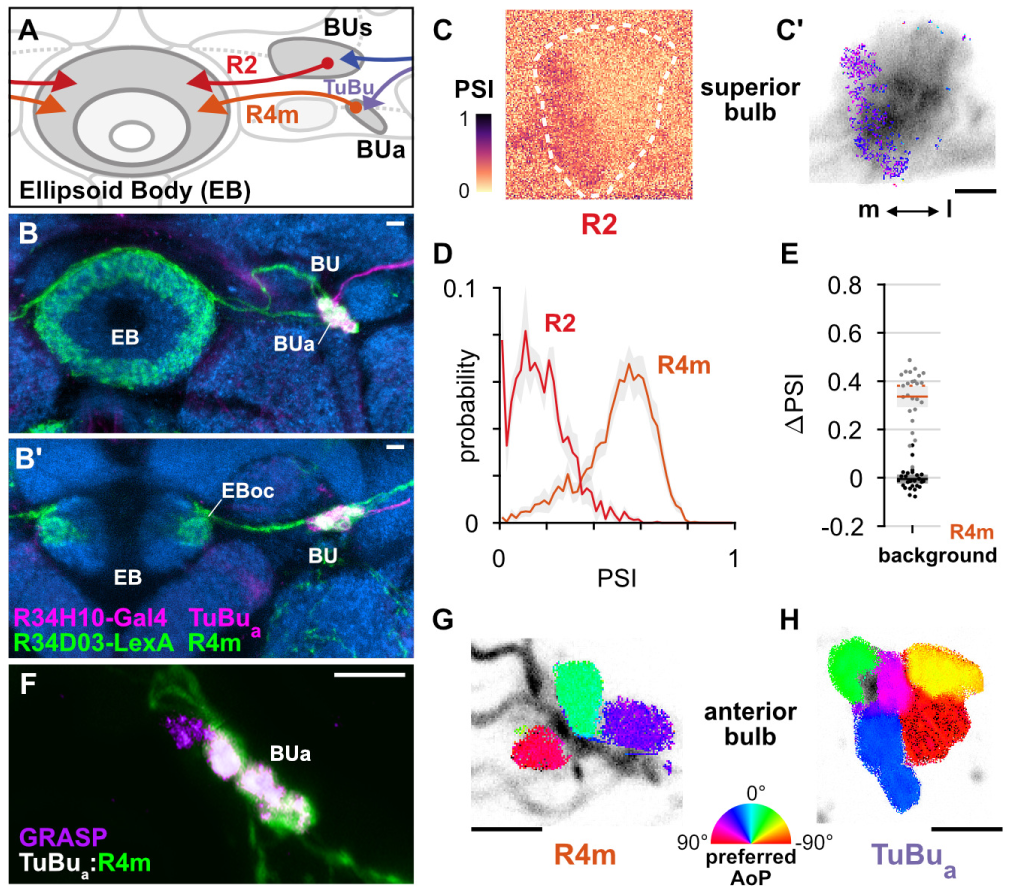
R4m ring neurons receive polarization-tuned responses from TuBu neurons. **A**: Schematic of TuBu and ring neuron connectivity in the bulb (BU). **B**: Confocal projection (anterior view) of dual-labeled TuBu_a_ neurons (R34H10-Gal4>RFP, magenta) and R4m neurons (R34D03-Gal4>GFP, green). **B’**: Dorsal view. Scale bars denote 5 μm. **C**: Example spatial maps of polarization-selectivity index (PSI) and tuning (**C’**) for R2 dendrites recorded in the superior bulb (R19C08-Gal4>GCaMP6s). Scale bar denotes 5 μm. **D**: Probability distributions of PSI values in ring neurons recorded in the bulb (average PSI **R2** neurons: 0.17, CI 0.05, background: 0.20, CI 0.03, N = 4 animals, p = 0.29 t-test; **R4m** neurons: 0.51, CI 0.11, background: 0.22, CI 0.05, N = 25 animals, p < 10^−6^ t-test). Mean ± CI. **E**: Effect of polarizer on median PSI values versus controls with polarizer removed, within R4m neurons (light dots) and background regions (dark dots) in individual animals (mean ΔPSI **R4m** neurons: 0.34, CI 0.11, N = 25, p < 10^−6^ t-test, background: −0.05, CI 0.05, N = 25, p = 0.58, t-test). **F**: Confocal projection (anterior view) of activity-dependent synaptic GRASP (GFP reconstitution across synaptic partners) signal between presynaptic TuBu_a_ and postsynaptic R4m neurons in the anterior bulb (BUa) (Macpherson et al., 2015). Scale bar denotes 5 μm. **G**: Example polarization tuning map in R4m dendrites in BUa (R34D03-Gal4>GCaMP6s). Pixels falling outside an ROI drawn around the neurons of interest, show average intensity in grayscale. Individual axons projecting medially to the EB are visible leaving the left side of the image. Scale bar denotes 5 μm. **H**: As in **G**, for TuBu_a_ output micro-glomeruli at an approximately corresponding location in BUa (R34H10-Gal4>sytGCaMP6s).

As with TuBu_s_ micro-glomerular outputs, we found that only a subset of R2 neurons in the BUs were modulated by polarized light, with above-threshold PSI values typically in a medial cluster with a preferred angle of polarization (AoP) around 45° (Fig. 6C). Low PSI values were common throughout the R2 population and average values were not significantly different from average values in background regions (Fig. 6D). By contrast, in R4m neurons in the BUa, average PSI values were greater than 0.5 and the overall distribution of values in the population was similar in shape to the distribution in TuBu_a_ neurons (Fig. 4B, Fig. 5B, Fig. 6D). We found that the polarizer had a significant effect on PSI values of R4m neurons versus controls with the polarizer removed (Fig. 6E). Furthermore, we found that the dendrites of individual R4m neurons exhibited distinct preferences for AoP in each recording (Fig. 6G). Since R4m neurons appear to receive monosynaptic input from TuBu neurons, we conclude that they almost certainly acquire their polarization-tuned responses from the presynaptic TuBu_a_ neurons in the BUa (Fig. 6A,B,F). We note that the average PSI value decreased from TuBu_a_ neurons to R4m neurons (Fig. S5) and we further explore the transformation of their signals in the next section. Although the BUs appears to contain polarization-sensitive elements, they are pervasive neither in the populations of R2 neurons nor their putative presynaptic partners, TuBu_s_ neurons, and hereafter we focus on polarization processing in the BUa.

In contrast to the linear polarotopic organization of tunings observed in the AOTU, which was consistent across animals (Fig. 4F,H), the spatial organization of polarization tunings in the BUa was less clear (Fig. 6G,H). We tested whether there was a common relationship between the horizontal (medial-lateral), vertical (ventral-dorsal), or angular position of micro-glomeruli within the BUa and their preferred AoP, for both TuBu_a_ and R4m neurons (Fig. S5). We also considered whether there was a relationship within a population of neurons in an individual animal which was not common across animals. We found no indication of a relationship between position and preferred AoP except in recordings of TuBu_a_ neurons in the left BUa, which showed a common vertically organized polarotopy (Fig. S5B) and circularly organized polarotopies in individual animals (Fig. S5C). However, we found no significant polarotopy in the corresponding TuBu_a_ neurons in the right BUa, or in postsynaptic R4m neurons. Hence we cannot firmly conclude that either a vertical or circular organization of tunings exists in the anterior bulb. Furthermore, our assessment of circular organization is only valid for the dorso-posterior imaging plane used here, and we cannot exclude the possibility of a circular organization around a different axis of the bulb.

### Populations of R4m ring neurons exhibit a preferred angle of polarization

We next wanted to understand how polarized light signals are represented in the ellipsoid body (EB), where the tangential ring neurons supply visual information around its circular structure. Ring neurons interact bidirectionally with columnar neurons (Omoto et al., 2018), which have been shown to flexibly encode heading direction relative to visual landmarks (Fisher et al., 2019; Kim et al., 2019; Seelig and Jayaraman, 2015). We recorded the synaptic terminals of the population of R4m neurons in the EB (approximately ten neurons, five per brain hemisphere, R34H10-Gal4). As expected from recordings in the dendritic regions of R4m in the anterior bulb (BUa), we observed modulation of their activity with rotations of the polarizer, indicated by their polarization-selectivity index (PSI) (Fig. 7A). Individual terminals were found to exhibit distinct tunings, and a range of tunings could be found intermingled at any given position in the EB (Fig. 7B). We noted here that in some recordings, above-threshold PSI values were spatially localized to approximately one quadrant of the EB (Fig. 7A,B, top, arrowhead). Additionally, we found that in many recordings the preferred angles of polarization (AoPs) of terminals were similar to each other within a recording, and the range of AoPs varied across animals (Fig. 7B). Therefore, the frequency of preferred AoPs was a unimodal distribution centered on a different angle in each recording (Fig. 7C). We verified that the non-uniform distribution of AoPs was not an artifact of our image projection across multiple planes and that a predominant preferred AoP was also observed from a single imaging plane through a section of the EB (Fig. 7A–C, bottom). As a result of these non-uniform tuning distributions, it followed that the average activity of the entire R4m population in the EB exhibited modulation induced by the polarizer and a single preferred AoP could effectively be identified for the population (Fig. 7D).

**Figure 7:**
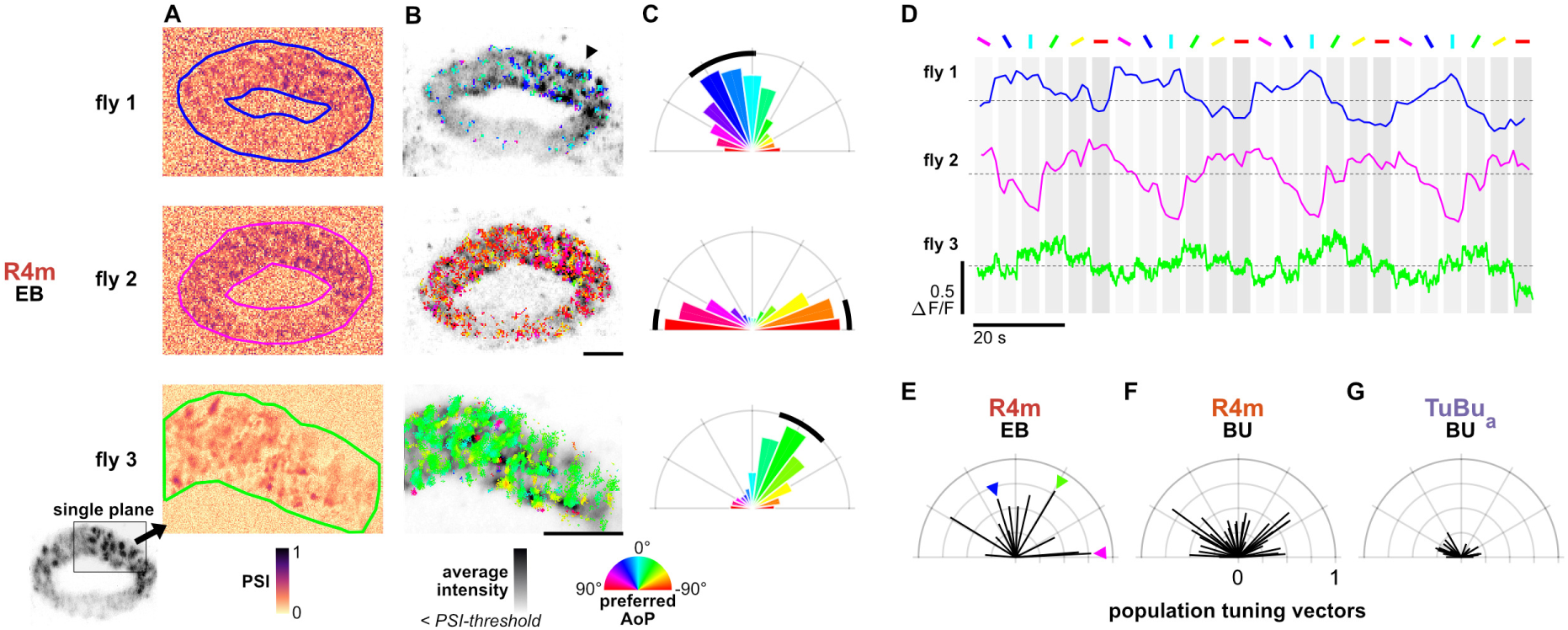
Populations of R4m ring neurons exhibit a preferred angle of polarization. **A**: Example spatial maps of polarization-selectivity index (PSI) in R4m synapses recorded in the ellipsoid body (EB) (R34D03-Gal4>sytGCaMP6s). Data shown are from maximum-selectivity projections through the EB (top, middle) or a single plane (bottom). **B**: Example polarization tuning maps corresponding to recordings in **A**. Pixels with a below-threshold PSI value, or falling outside an ROI drawn around the R4m population, show average intensity in grayscale. Scale bars denote 10 μm. **C**: Polar histograms of preferred angles of polarization in all pixels within the ROIs in **A**. Normalized probabilities in each bin are displayed as area of wedge; radial lengths of wedges not directly comparable. Arc denotes mean resultant angle ± 95% confidence interval (**fly 1**: 0.57 18.7° CI 16.6 °, N = 4, p = 0.002 Rayleigh uniformity test; **fly 2**: 0.72 −87.3° CI 15.0°, p = 0.001 Rayleigh uniformity test; **fly 3**: 0.71 −31.6° CI 15.4°, p = 0.001 Rayleigh uniformity test). **D**: Average GCaMP activity in the ROIs in **A** in response to different angles of polarization. **E**: Resultant tuning vectors for the population of all recorded R4m synapses in the EB of individual animals (mean length, pixel-based: 0.51, CI 0.44, N = 7, p < 10^−6^ t-test). Arrowheads indicate data for examples in **A-D**. **F**: Resultant tuning vectors for the population of all recorded R4m neurons recorded in the left or right BU of individual animals (mean length, pixel-based: 0.39, CI 0.32, N = 25, p < 10^−6^ tailed t-test; ROI-based: 0.36, CI 0.46, N = 25, p = 0.005 tailed t-test, 134 ROIs, > 3 ROIs per BU). **G**: Resultant tuning vectors for the population of all recorded TuBu_a_ neurons recorded in the left or right BU of individual animals (mean length, pixel-based: 0.18, CI 0.13, N = 7, p < 10^−6^ tailed t-test; ROI-based: 0.14, CI 0.15, N = 7, p = 0.0002 tailed t-test, 101 ROIs, > 3 ROIs per BU).

To compare the distribution of tunings across animals, we calculated the mean resultant vector of the tunings of all pixels within the EB, weighted by their individual PSI values (Fig. 7E). The length of the vector gives an indication of the distribution of polarization tunings in a single recording, with a value of 1 indicating an identical preferred AoP in all pixels and a value of zero indicating a uniform distribution of preferred AoPs. For R4m terminals in the EB we found population tuning vectors with lengths exceeding 0.74 and an average length of 0.51 across animals (Fig. 7E), while for R4m dendrites recorded in either the left or right BUa individually we found an average length of 0.39 (Fig. 7F). For TuBu_a_ populations recorded in either bulb we found that the vector lengths did not exceed 0.3 and the average length was 0.18 across animals (Fig. 7G). Since uneven sizes or quantities of neurons could affect these results, we repeated the analysis with ROIs drawn on individual micro-glomeruli in the bulb. We found a comparable number of micro-glomeruli in recordings of TuBu_a_ and R4m neurons in the BUa, and the ROI- and pixel-based approaches both yielded a qualitatively similar result (Fig. 7F,G).

These findings suggest that there is not an exact correlation between polarized light responses in the populations of presynaptic TuBu_a_ neurons and postsynaptic R4m neurons in an individual animal. In R4m dendrites, the average strength of modulation is reduced compared to TuBu_a_ neurons (Fig. S5) and the distribution of tunings is less uniform (Fig. 7F,G). In R4m terminals in the EB, the distribution of tunings is less uniform still, hinting at subcellular processes which may impact R4m signalling locally in the EB, a computational motif for which there is precedence both in the CX and in visual neurons generally (Franconville et al., 2018; Turner-Evans et al., 2020; Yang et al., 2016). As a consequence, it appears that the ensemble activity of R4m synapses could convey a preferential response for a particular angle of polarization to columnar neurons at any location in the EB.

### E-PG neurons respond to polarized light with flexible tuning and no fixed polarotopic map

We then asked whether columnar E-PG neurons (also referred to as ‘compass’ neurons) respond to polarized light cues. E-PG neurons are key elements in a network which maintains a neural representation of heading direction as a locus of activity, or ‘bump’, which changes position within the CX as the animal turns, like the needle of a compass (Green et al., 2017; Seelig and Jayaraman, 2015). In the previous literature, this activity bump has been observed in the ellipsoid body (EB), protocerebral bridge (PB), and fan-shaped body (FB), typically during walking or flight in restrained animals (Giraldo et al., 2018; Shiozaki et al., 2020). It has not been demonstrated in fully immobilized animals, hence we did not expect to see it here. Nevertheless, we hypothesized that E-PG activity could be modulated by a varying angle of polarized light since the same has been demonstrated in numerous columnar central complex neurons in other insects (Heinze and Homberg, 2007; Honkanen et al., 2019). Moreover, the responses we observed in R4m ring neurons (Fig. 7D) suggested that the E-PG population should also exhibit tunings to a limited range of angles. Ring neurons provide inhibitory input to E-PG neurons in the EB (Fig. 8A), where interactions between ring and E-PG neurons are thought to be reciprocal (Fisher et al., 2019; Kim et al., 2019; Omoto et al., 2018). Using activity-dependent GRASP (Macpherson et al., 2015), we found labeling of synapses between presynaptic E-PG neurons and postsynaptic R4m neurons in the EB (Fig. S6B), confirming the reciprocal connectivity between the neurons in the respective drivers (R4m: R34D03-LexA, Fig. 6B; E-PG: SS00096-Gal4, Fig. S6A).

**Figure 8:**
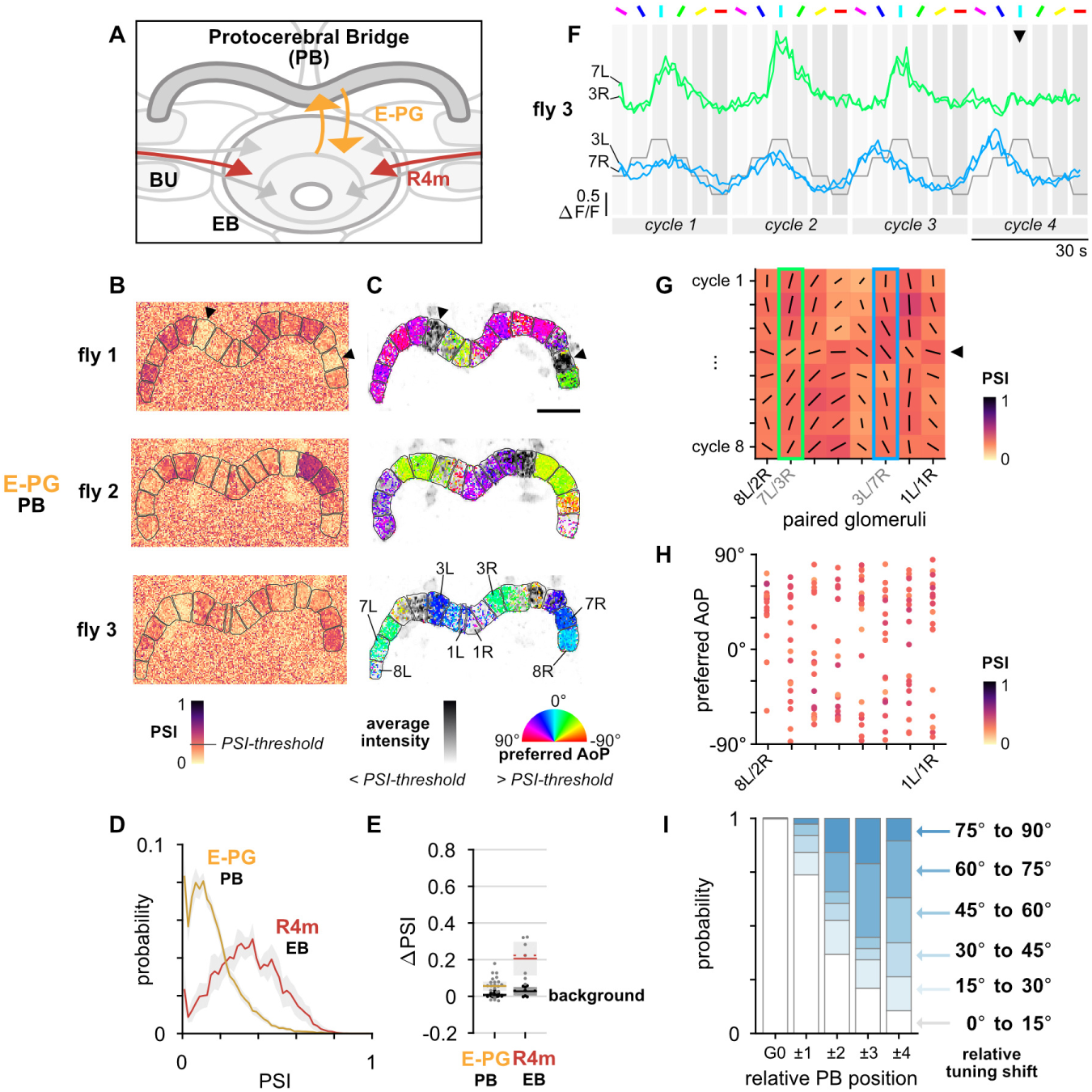
E-PG neurons respond to polarized light with flexible tuning and no fixed polarotopic map. **A**: Schematic of E-PG columnar neuron projections and connectivity with tangential ring neurons in the ellipsoid body (EB). See also Fig. S6E. **B**: Example spatial maps of polarization-selectivity index (PSI) in E-PG synapses recorded in the protocerebral bridge (PB) (SS00096-Gal4>sytGCaMP6s). Data shown are from maximum-selectivity projections through the PB. ROIs (gray) demarcate glomeruli. **C**: Example polarization tuning maps corresponding to recordings in **A**. Pixels with a below-threshold PSI value, or falling outside an ROI drawn around the PB, show average intensity in grayscale. Scale bar denotes 25 μm. **D**: Probability distributions of PSI values in E-PG neurons recorded in the PB and R4m neurons recorded in the EB (average PSI **E-PG** neurons: 0.14, CI 0.05, background: 0.19, CI 0.01, N = 22 animals, p = 0.0001 t-test; **R4m** neurons: 0.34, CI 0.11, background: 0.21, CI 0.03, N = 7 animals, p = 0.02 t-test). Mean ± CI. **E**: Effect of polarizer on median PSI values versus controls with polarizer removed, within E-PG and R4m neurons (light dots) and background regions (dark dots) in individual animals (mean ΔPSI **E-PG** neurons: 0.06, CI 0.05, N = 22, p < 10^−4^ t-test, background: 0.01, CI 0.01, N = 22, p = 0.0007, t-test; **R4m** neurons: 0.21, CI 0.11, N = 7, p = 0.002 t-test, background: 0.03, CI 0.03, N = 7, p = 0.04, t-test). **F**: Activity in two pairs of L/R ROIs in **C** (fly 3) in response to different angles of polarization. Arrowhead indicates position of expected peak. **G**: Cycle-by-cycle characterization of E-PG responses across the PB in a single recording (fly 3, **C**). Vector orientation represents preferred AoP, length represents PSI (grid spacing equal to 1). Highlighted boxes indicate extended data for pairs shown in **F**. Arrowhead indicates the same cycle as the arrowhead in **F**. **H**: Scatter plot showing position of paired E-PG glomeruli in the PB and preferred angle of polarization (AoP) (pooled data ρ = 0.23, N = 19 animals, p = 0.006 permutation test, 152 ROIs, mean ROI PSI 0.34 ± 0.06; 5 significant individual circular-circular correlations, mean ρ = 0.46, SEM 0.45). **I**: Normalized probability of tuning shift magnitude with distance from the glomerulus with the highest PSI value (mean shift between positions 2 to 3, p = 0.21; 3 to 4, p = 0.65; 2 to 4, p = 0.08; all other pairs p < 10^−3^, N = 19 animals, 152 ROIs). See also Fig. S6G.

We then recorded calcium signals in the presynaptic terminals of E-PG neurons in the PB, where they form 16 distinct glomeruli (Fig. 8A), each innervated by at least two E-PG neurons (Fig. S6) (Wolff et al., 2015). Due to their neighboring positions in the EB and connectivity with other neurons, the activity of E-PG neurons innervating the 8 glomeruli in the left half of the PB is known to be coordinated with those in the 8 glomeruli in the right half (Fig. S6E), and on either side of the PB the ends are effectively wrapped (1L is continuous with 8L, 1R is continuous with 8R) (Giraldo et al., 2018; Green et al., 2017). We found that E-PG activity in the PB was modulated as the polarizer was rotated. We assigned PSI values to the pixels in each recording as an indicator of modulation (Fig. 8B) and calculated their preferred angle of polarization (AoP) (Fig. 8C). As expected, the PSI values and preferred AoPs showed a bilateral coupling, with the right half of the PB (1R to 8R) resembling the left half (8L to 1L) (Fig. 8B,C). In different animals, the preferred AoP varied in glomeruli at corresponding positions in the PB (Fig. 8C). We also observed that the distribution of PSI values was not homogenous across the PB, and high values typically clustered across a contiguous subset of 2–4 glomeruli, while low PSI values occurred throughout the remaining glomeruli (Fig. 8B). Across the glomeruli in each cluster, the preferred AoP was similar in a given animal (Fig. 8C). It should be noted that these clusters of high PSI values correspond to the regions of highest modulation over a period of minutes, not an instantaneous locus of intensity which moved across the PB (activity bump) (Giraldo et al., 2018; Green et al., 2017). Indeed, glomeruli with high average intensities often exhibited low PSI values (arrowhead, Fig. 8B,C).

Overall, we found substantially lower PSI values in E-PG neurons than in R4m neurons (Fig. 8D). We found a statistically significant effect of the polarizer on PSI values versus controls in both populations (Fig. 8E), yet in E-PG neurons the effect size was small and the average PSI value was generally lower than in background regions of recordings (Fig. S6C). To explore this discrepancy, we examined the responses of individual glomeruli in the PB in response to cycles of the polarizer (Fig. 8F). Here, in the PB, we observed characteristics which distinguished the responses from those of all other polarization-sensitive elements that we recorded in the upstream pathway. First, the amplitude of responses was often found to be inconsistent over multiple rotation cycles of the polarizer (Fig. 8F, top). Second, the peak response was often found to occur at different positions of the polarizer over multiple cycles (Fig. 8F, bottom). For both of these response characteristics, variations were synchronized across the left and right PB glomerulus pair (Fig. 8F). When we analyzed responses to individual cycles of the polarizer separately, these characteristics manifested as PSI values and preferred AoPs which varied over time (Fig. 8G). To obtain a measure of synchronicity between E-PG modulation and the polarizer stimulus, we examined the auto-correlation function of all individual glomerular responses, and compared them with those of R4m and TuBu_a_ neurons recorded in the anterior bulb (BUa). For E-PG neurons, we found that less than half of all glomeruli recorded exhibited a periodicity which matched the stimulus, while almost all R4m and TuBu_a_ neurons matched the stimulus (E-PG: 43.3%, R4m: 98.4%, TuBu_a_: 100%) (Fig. S6D). Therefore, although periodic, when observed over multiple cycles the majority of E-PG responses were found to be no more synchronized with the rotation of the stimulus than the fluctuations in their activity recorded with the polarizer removed (Fig. S6D). This finding is reminiscent of the observation of ‘conditional’ polarization-sensitivity in some columnar neuron types in the locust central complex (Heinze and Homberg, 2009). While we did not specifically test the stability of R4m responses recorded in the EB as we could not distinguish individual neurons, it should also be noted that the E-PG activity analyzed also potentially represents multiple neurons per glomerulus which could have been differentially active. Nevertheless, their activity profiles (Fig. 7D, Fig. 8F) and the difference in their average PSI values (Fig. 8D,E) indicate that, if E-PG polarization-sensitivity does indeed result from R4m input, an additional transformation of signals occurs between these neurons.

We next sought to address the organization of preferential responses to polarized light in the PB, acknowledging that neither the preferred angles of polarization nor the PSI values calculated for E-PG neurons were necessarily stable over time (Fig. 8G). We therefore limited our analysis to individual cycles of the stimulus, and we pooled the coordinated responses of glomeruli from the left and right sides of the PB. To evaluate the most appropriate pooling, we cross-correlated the activity recorded from pairs of left and right glomeruli under different pairing schemes and found the normalized coefficient as an indication of their similarity (Fig. S6E). The pairing scheme following the logic 1L/1R, 8L/2R, 7L/3R, etc. (Fig. S6E) yielded the highest mean similarity across all glomeruli, which decreased with a sinusoidal profile as the distance between pairs increased (Fig. S6F). This pairing confirms a scheme proposed based on anatomical connectivity (Wolff et al., 2015), but differs by one position from the proposed connectivity in the locust, where a pairing scheme corresponding to 1L/8R, 8L/1R, 7L/2R, etc. (Fig. S6E) has previously been used to pool data (Heinze and Homberg, 2009).

Across animals, we found no common relationship between glomerulus position in the PB and the preferred angle of polarization (AoP) of E-PG neurons (Fig. 8H), matching the findings for the homologous CL1a neurons in locusts (Heinze and Homberg, 2009; Pegel et al., 2019). We then asked whether, on the timescale of a single stimulus cycle (30 s), there was any relationship between PB position and preferred AoP in an individual animal. In each recording, we picked at random a single response cycle in which the average PSI value across all glomerulus pairs exceeded a threshold (mean + 1 SD of PSI values in background regions of all E-PG recordings). We then identified the glomerulus pair with the maximum average PSI value, which we refer to as G0, and expressed all preferred AoPs, PSI values and positions in the PB relative to G0 (Fig. S6G). Smooth transitions in preferred AoP across glomeruli were observed infrequently, and in 6 out of 19 animals this resulted in a weak relationship between PB position and preferred angle of polarization (asterisks, Fig. S6G).

More generally, we found that glomeruli neighboring G0, at ± 1 PB position, were likely to exhibit a similar preferred AoP to G0, to within 15° (Fig. 8I, Fig. S6G). At ± 2–4 PB positions from G0, we found preferred AoPs generally shifted towards orthogonal angles (Fig. 8I, Fig. S6G) and among these positions there was again a similarity between neighboring glomeruli (Fig. S6H). These data support our initial observation of clusters of glomeruli with similar tunings and PSI values (Fig. 8B,C), contrasting with the polarotopic organization of tunings across the PB found for CPU1 neurons in locusts (likely homologous to P-F-R neurons in flies) (Heinze and Homberg, 2007; Honkanen et al., 2019; Pegel et al., 2019). A limited representation of two orthogonal angles of polarization in columnar neurons would also be congruent with a single predominant tuning being conveyed by the R4m population (Fig. 7D), since rectification of a sinusoidal tuning function would directly lead to two signals with peak responses at orthogonal angles.

## DISCUSSION

In this study we have demonstrated that each section of the *Drosophila* anterior visual pathway (AVP) contains polarization-tuned neurons. Together, they provide a circuit to convey polarized light signals from the specialized dorsal rim area of the eye to the compass neurons of the central complex, via the anterior optic tubercle and bulb. This pathway also conveys information about unpolarized visual features, as shown here and in previous studies. The encoding of multiple visual modalities, the similarities in the constituent neurons, and the organization of the neuropils which accommodate them (Omoto et al., 2017), support the view that the AVP in *Drosophila* is homologous to the sky compass pathway described in locusts, bees, butterflies, and beetles, among other insects (Honkanen et al., 2019; Warren et al., 2019).

Our approach to investigating the neural processing of polarization vision offered a number of advantages over traditional intracellular electrophysiology. Firstly, it allowed us to simultaneously record from whole populations of neurons, which would otherwise be technically challenging. Here, we exploited this to investigate the spatial organization of polarization responses in an individual animal. This may be key in understanding the central complex, where dynamic responses reflect circuit plasticity and depend on numerous factors, such as proprioceptive inputs, internal states and goal-direction. Next, targeted expression of calcium indicators allowed us to isolate specific anatomical groups of neurons, such as specific TuBu or ring neuron populations, greatly increasing the repeatability of functional characterizations. Crucially, the identification of corresponding genetic drivers will enable silencing experiments, optogenetic stimulation and multi-population recordings to probe circuit function in the future. Imaging of calcium indicators also facilitated the characterization of neurons whose axons are prohibitively thin for recording intracellularly. MeTu-like neurons, for example, have long been assumed to deliver polarization signals from the medulla to the anterior optic tubercle, and here we were able to confirm this by direct observation for the first time.

### Skylight polarization features extracted by the MEDRA

Since each detector for polarized light in the DRA essentially has a different field of view, the success of this approach depended on the ability to stimulate a sizable number of DRA ommatidia. Surprisingly, almost the full extent of the DRA was stimulated by polarized light originating from a single point in the visual field with a common angle of polarization. A wide range of polarization tunings was subsequently revealed in downstream neurons, supporting the idea that the *Drosophila* medulla dorsal rim area (MEDRA) analyzes the overall pattern of polarized light in the sky and extracts a predominant angle of polarization (AoP) (Labhart, 2016; Rossel and Wehner, 1986), rather than performing many local AoP estimates. During the morning and evening when *D. melanogaster* are most active, the pattern of polarization in the sky can be well approximated by a single, predominant AoP. DmDRA1 neurons appear to spatially integrate polarization signals from multiple columns of the MEDRA (Fig. 1), and individual neurons heavily overlap each other (Sancer et al., 2019). This could provide an additional robustness to occlusions of the sky or of the DRA itself and average out inconsistencies in the available light (Labhart et al., 2001; Rossel and Wehner, 1986).

The parallel circuitry between DRA R7, DmDRA1 and MeTu neurons in MEDRA columns (Fig. 2D), resembles the color-processing pathway found in non-DRA columns involving R7, Dm8 and Tm5c (Gao et al., 2008; Karuppudurai et al., 2014). MeTu neurons in the MEDRA may also integrate color signals, as their dendritic fields extend into the non-DRA medulla, indicating that color and polarization processing are compatible (Fig. S3). We have not functionally described the responses of DmDRA2 cells that contact R8 cells in this study (Sancer et al., 2019), and these cells may be differently integrated with color processing. Both parallel functions will likely need to be incorporated to build a complete conceptual model of skylight polarization processing in the medulla.

### Sensory transformations through the AVP

In the anterior optic tubercle (AOTU), we found polarization-sensitive neuron populations entering and leaving the tubercle via the intermediate-lateral domain (Fig. 2–4). We also observed polarization responses in the lateral domain, although it is unclear whether this is a result of separate polarization-sensitive MeTu types projecting from the MEDRA to different AOTU domains. Alternatively, since MeTu neurons are also postsynaptic in the AOTU (Omoto et al., 2017), signals from a single polarization input channel could be redistributed to different regions of the AOTU for integration with other visual modalities or bilateral interactions (Fig. 3). The AOTU in *Drosophila* is also likely to be a site for modulation of signals depending on time or internal states (Guo et al., 2018; el Jundi et al., 2014; Lamaze et al., 2018), and a capacity to modify responses may explain why we observed multiple polarotopic organizations in a MeTu neuron population in the AOTU (Fig. S3). However, there may also be multiple functional subtypes within the population that more tailored experiments may be able to distinguish.

Intriguingly, none of the polarotopies found in presynaptic MeTu neurons (Fig. 2L,M) matched the polarotopy of postsynaptic TuBu dendrites in the AOTU (Fig. 4G,I), which was extremely consistent across animals. Our findings suggest that TuBu neurons extract a processed form of the signals in the AOTU, encoding visual features within fewer neurons than the MeTu populations. TuBu neurons appear to divide signals into functional groups, and the anterior bulb-projecting TuBu_a_ group in every fly contained a set of around six tunings covering −90° to +90° of polarization space in approximately 30° steps, tightly-packed in a micro-glomerular structure with no apparent polarotopy (Fig. 5, Fig. 6). The question remains open as to whether a sun position system and skylight polarization system are independent in the bulb. Unlike the TuLAL neurons in locusts (homologous to TuBu), where there is convergence on the dendrites of postsynaptic neurons (Hadeln et al., 2020; Pegel et al., 2018; Pfeiffer et al., 2005), TuBu neurons appear to form one-to-one contact with individual ring neurons (Omoto et al., 2017). Hence, we posit that the site of integration of celestial cues is not at the synapse between TuBu and ring neurons. Although we found evidence that angles of polarization are represented in the superior bulb (Fig. 5, Fig. 6), where unpolarized cues are also known to be represented, the populations we recorded contained a limited range of tunings and resembled a system for detecting visual features with a particular polarization signature (Labhart, 2016), such as horizontally polarized light reflected from surfaces like water, rather than a system for accurate estimation of orientation. Such responses would likely be mediated by more ventral regions of the eye than the DRA (Velez et al., 2014; Wernet et al., 2012). It should be noted that our polarized light stimulus broadly illuminated the eye from a dorsal position and, although we attempted to minimize reflections, we did not measure whether reflected polarized light fell on the ventral eye during our experiments.

### Stereotypic polarotopy in the periphery gives way to idiosyncratic plasticity in the CX

By recording the ensemble response of a population of R4m ring neurons, both in the anterior bulb and ellipsoid body (EB), we determined that they do not simply relay the responses of presynaptic TuBu_a_ neurons to the EB. Instead, they appear to deliver a subset of signals more prominently than others, bestowing the population with an ensemble response tuned to a specific angle of polarization (Fig. 7). Furthermore, we found that this population tuning conveys a different angle of polarization in individual animals, and one exciting possibility is that this represents a flexible heading signal relative to polarized light cues, which could direct behavior (Warren et al., 2018). A question to address in future work is whether the preferred angle of polarization of an individual ring neuron is itself fixed, in which case we may have observed the result of a winner-take-all competition among the R4m population in the EB, or if the whole population flexibly re-tunes to preferentially respond to a common AoP. Recordings from individual neurons will be required to resolve this.

It is clear that among R4m and E-PG neurons, polarization tunings are not represented with a retinotopic map in the EB or PB which is common between individual animals (Fig. 7, Fig. 8). This is in contrast with the consistent polarotopic organizations found upstream in the MEDRA or AOTU (Fig. 1–4), but in agreement with a previous study which showed that the azimuthal position of unpolarized visual stimuli is also not represented retinotopically in E-PG neurons (Fisher et al., 2019). The lack of organization in E-PG responses also matches previous findings in the corresponding CL1a neurons in locusts, but contrasts with the polarotopic organization found in other columnar neurons in the locust CX, such as CPU1, and the tangential TB1 neurons (Heinze and Homberg, 2007, 2009; Pegel et al., 2019). A potential explanation for the lack of consistent polarotopy in CL1a, or indeed E-PG neurons, was offered by Heinze and Homberg (2009): at least two of each neuron type innervates an individual glomerulus in the PB. Could each of these have differential responses to polarized light to enable different configurations across the PB? Intriguingly, the TB1-like Δ7 neurons in the *Drosophila* PB appear to synapse onto only a subset of the E-PG neurons in a single glomerulus (Turner-Evans et al., 2020), perhaps indicating independent functional groups. We may therefore yet find a polarotopic organization of responses in the *Drosophila* CX. Alternatively, such an organization may reflect a common, genetically pre-programmed directional goal to facilitate migration, which flies may lack (Honkanen et al., 2019), instead using polarization cues to follow a fixed course and disperse along idiosyncratic headings (Dickinson, 2014).

Our data suggest that in a given fly, E-PG neurons may respond to one of two approximately orthogonal angles of polarization, effectively dividing the population into two groups. Interestingly, when data from locust CPU1 neurons (likely homologues of P-F-R neurons in *Drosophila*) were pooled with tunings obtained from a number of other polarization-sensitive columnar CX neuron types, including CL1b (P-EG), CL2 (P-EN), CPU2, and CPU4 (P-FN), the organization of tunings in the locust PB could be interpreted as clustering around two orthogonal preferred angles (Heinze and Homberg, 2009). A binary system such as this would be well suited to influence downstream processes in a motor-centered coordinate frame (Rayshubskiy et al., 2020). For example, the eventual output of the compass network may be a command signal to activate one descending neuron of a bilateral pair to initiate a turn to either the left or right, and thus maintain a heading specified by polarization patterns in the sky.

An important next step will be to understand how polarized light influences the activity bump in columnar neurons and whether the activity of columnar neurons reciprocally influences the tunings of R4m neurons. We did not observe an activity bump in E-PG neurons in the PB, likely due to the open-loop stimulus presentation and recordings performed in immobilized animals, although we could see evidence of flexible encoding of polarization information (Fig. 8). According to our mappings of E-PG responses in the PB, the influence of a rotating polarized light stimulus might be to move the activity bump discontinuously between two positions, not dissimilar to observations in a recent investigation of the influence of airflow on the bump in E-PG neurons (Okubo et al., 2020). However, a limitation of the polarization stimulus used here is that the intensity gradient and position of the light source did not change as the angle of polarization rotated, as it would be seen to by an animal turning under a natural sky. If the ambiguity between 0/180° polarization cues is resolved by integrating light intensity information, then the stimulus we used here presented contradictory, unnatural changes. Behavioral studies in ants (Wehner and Müller, 2006) and dung beetles (el Jundi et al., 2015) have demonstrated that skylight polarization cues can have a greater influence than other visual features in guidance and navigation behaviors, while in *Drosophila* intensity gradients appear to have a greater behavioral significance (Warren et al., 2018). A key challenge for future studies will be to uncover the mechanisms for integrating and selecting from the multiple sensory modalities and visual qualities represented in the central complex in order to navigate complex environments.

## Acknowledgments

We are grateful to Sam LoCascio for technical advice. Tanya Wolff and Vivek Jayaraman kindly provided the split-Gal4 line SS00096. We also thank Holger Krapp, Kit Longden, and members of the Frye lab for their comments on the manuscript. Stocks obtained from the Bloomington Drosophila Stock Center (NIH P40OD018537) were used in this study. This work was supported by grants from the NIH (R01-NS096290 to V.H. and R01-EY026031 to M.A.F.).

## Author contributions

Ordered according to main list of authors:

**Conceptualization**: B.J.H., J.J.O., V.H., M.A.F.

**Data curation**: B.J.H., P.K., B.-C.M.N.

**Formal analysis**: B.J.H., J.J.O., P.K., B.-C.M.N.

**Funding acquisition, resources, administration**: V.H., M.A.F.

**Investigation**: B.J.H., J.J.O., P.K., B.-C.M.N., M.F.K., N.K.B.

**Methodology**: B.J.H., J.J.O., M.F.K.

**Software, validation**: B.J.H.

**Supervision**: B.J.H., J.J.O., V.H., M.A.F.

**Visualization**: B.J.H., J.J.O., P.K., V.H.

**Writing – original draft**: B.J.H.

**Writing – review & editing**: B.J.H., J.J.O., P.K., V.H., M.A.F.

## METHODS

### In vivo calcium imaging

#### Fly preparation

Flies were raised at 25°C on a standard cornmeal/molasses diet in 40 ml vials, under a 12:12 hour dark:light cycle. Imaging experiments were performed between ZT0–14, although time of day was not a factor in our experimental design or analysis. We imaged 1–7 day old female flies expressing either UAS-GCaMP6s (Chen et al., 2013) for dendritic regions or UAS-sytGCaMP6s (Cohn et al., 2015) for axon terminals, together with UAS-tdTomato (Shaner et al., 2004) for image registration. Flies were cold anaesthetized and mounted on a custom fly holder, modified from (Weir et al., 2016), with the head pitched forward so that its posterior surface was approximately horizontal (Fig. S1A). Surfaces of the fly holder visible to the fly were covered in matte white paint (Citadel) and roughened to reduce confounding reflected polarized light cues (Foster et al., 2018). We fixed the fly to the holder using UV-curing glue (Fotoplast) around the posterior-dorsal cuticle of the head and at the base of the wings on either side of the thorax. To reduce movement of the brain we fixed the legs, abdomen and proboscis with beeswax. We used forceps to remove the cuticle and air-sacs above the optic lobe or central brain, depending on the recording site, and cut muscle 1 (Demerec, 1950) to reduce movement. Physiological saline (103 mM NaCl, 3 mM KCl, 1.5 mM CaCl_2_, 4 mM MgCl_2_, 26 mM NaHCO_3_, 1 mM NaH_2_PO_4_, 10 mM trehalose, 10 mM glucose, 5 mM TES, 2 mM sucrose) was perfused continuously over the brain at 1.5 ml/min via a gravity drip system and the bath was maintained at 22°C for the duration of experiments by an inline solution heater/cooler (SC-20, Warner Instruments) connected to a temperature controller (TC-324, Warner Instruments).

#### Imaging setup

We used a two-photon excitation scanning microscope controlled by Slidebook (ver. 6, 3i) with a Ti:sapphire laser (Chameleon Vision, Coherent) at 920 nm and a 40× objective (0.8 numerical aperture, NIR Apo, Nikon). For each brain area imaged, we aimed to capture the full extent of the volume of labeled neurons, using a maximum step-size of 4 μm between imaging planes, and maintained a volume-rate of at least 1 Hz. Image resolution varied depending on the number of planes captured but was not less than 100 pixels in the longest dimension. We recorded frame capture markers and stimulus events on a DAQ (6259, NI) sampling at 10 kHz.

#### Polarized light stimulus

We used a custom polarized light stimulus device comprising a UV LED (M340D3, Thorlabs), a 7.5 mm diameter aperture, a ground glass diffuser (DGUV10-1500, Thorlabs), a low-pass filter (FGUV11, Thorlabs), and a removable linear polarizer (BVO UV, Bolder Optic). The UV LED was controlled through MATLAB 2017a (Mathworks, MA) via a DAQ (6259, NI) and LED driver (LEDD1B, Thorlabs). The polarizer was rotated with a bipolar stepper motor (ROB-10551, SparkFun) and spur gears (1:1), and a motor driver (ROB-12779, SparkFun) controlled through MATLAB (2017a, Mathworks) via a DAQ (USB1208, MCC), with a minimum step-size of 7.5°. The motor was operated in open-loop and a Hall effect sensor (A1324, Allegro) was used to detect the proximity of a magnet which passed once per revolution, in order to verify correct operation. Angles of polarization and directions of rotation are expressed from an external viewpoint looking towards the fly (Fig. S1A). 0°/180° corresponds to a vertical orientation in the transverse plane and an alignment with the fly’s long-axis in the horizontal plane. We investigated the reproducibility of the polarizer’s angular positions and measured <1° variation over multiple revolutions and <1° of position hysteresis (backlash) after reversing the direction of rotation. The surface of the polarizer was positioned frontally, 110 mm from the fly’s head at an elevation of approximately 65° above the eye-equator (Fig. S1A). The light subtended a solid angle of approximately 4° and the entirety of the fly, including the dorsal rim area of both eyes, was illuminated. We measured approximately 0.8 μW/cm^2^ irradiance at the fly’s head at the spectral peak of 342 nm (8.7 nm FWHM) with the polarizer attached (Fig. S1B). We calibrated the LED power in order to maintain a similar irradiance value with the polarizer removed (Fig. S1B). We measured a ± 5% modulation in light intensity over a full revolution of the device (Fig. S1B), due to a slight off-axis tilt of the diffuser and polarizer. This intensity modulation was of similar magnitude both with the polarizer attached and removed, and was therefore unlikely to be an effect of polarization. We reasoned that if calcium activity in neurons was modulated by the rotation of the device with the polarizer attached, but not with the polarizer removed, then the varying angle of polarization throughout the revolution was its cause, rather than the varying light intensity. To quantify the difference in modulation between these two polarizer conditions, we report the change in polarization-selectivity index (ΔPSI) throughout (see *Polarization-selectivity index*).

We verified that the polarized light stimulus elicited an expected response in the dorsal rim photoreceptors by recording calcium signals in R7/R8 terminals in the medulla dorsal rim area (MEDRA) (Fig. S1C–E). We observed preferential responses to different angles of polarized light across the MEDRA and approximately orthogonal preferred angles within R7/R8 pairs in individual columns (Fig. S1C–E). Moving anterior to posterior across the right MEDRA, the preferred angle of polarization rotated counter-clockwise (Fig. S1E), matching a previous characterization (Weir et al., 2016). We estimated that at least 80% of MEDRA columns were stimulated and conveyed polarization tunings that matched predictions based on the anatomy of photoreceptors at corresponding positions (Weir et al., 2016) (Fig. S1E–G), with weak responses or deviations observed only in the anterior-most columns (Fig. S1E,F) likely due to their posterior receptive fields which faced away from the stimulus. With the polarizer removed, we observed no spatial organization of tunings in photoreceptor terminals and PSI values close to zero (Fig. S1J), indicating reduced modulation of activity by the stimulus.

#### LED display

We used a 32 × 96 pixel display, composed of 8 × 8 panels of LEDs (470 nm, Adafruit) with controllers (Reiser and Dickinson, 2008), arranged in a half-cylinder spanning ± 90° azimuth from visual midline and approximately ± 30° elevation from the eye-equator (Fig. S1A). Each LED pixel subtended a solid angle of approximately 1.5° at the eye-equator. At their maximum intensity, we measured approximately 0.11 μW/m^2^ irradiance at the fly’s head at the spectral peak of 460 nm (243 nm FWHM).

### Experimental protocols

Visual stimuli were presented in sets as described below. Between each stimulus set, 10 s of spontaneous activity was recorded in darkness with no visual stimulation. The polarizer could only be removed or attached between recordings, but could be done so while maintaining the same imaging parameters and field-of-view under both conditions.

#### Angle of polarization tuning

To characterize responses to different angles of polarization, we rotated the polarizer discontinuously in 30° steps with the UV LED on throughout. Each of the 12 positions (6 unique angles of polarization) was maintained for 4–4.5 s and we used 4 s of imaging data collected during this period in our analysis. The polarizer was then rotated through 30° in 0.5 s. At least two complete revolutions of the polarizer were made. For recordings with the polarizer removed, the procedure was repeated and one revolution of the stimulus was made.

#### Polarized light flash

To characterize responses to individual wide-field flashes of polarized light, the polarizer was first rotated to 0° (vertical) in darkness. A series of three flashes of the UV LED were presented, 4 s on:4 s off. After 10 s the same procedure was repeated with the polarizer at 90° (horizontal). The light was the same used in the tuning protocol. For recordings with the polarizer removed, the procedure was repeated with flashes at the 0° position.

#### Unpolarized light flash

To characterize responses to individual wide-field flashes of unpolarized light, the entire LED display was illuminated following the same procedure as for polarized light flashes.

#### Bars

To characterize retinotopic responses to unpolarized stimuli, a single bright, vertical bar was presented on the LED display (32 × 1 pixel) with all other LEDs off (0.78 Weber contrast). Bars initially remained stationary for 3 s, then jittered left and right (± 1 pixel) for 3 s, followed by an inter-trial period of 4 s with all LEDs off. Bars were presented at five equally spaced azimuth positions spanning ± 90°, presented sequentially from left to right around the fly. This procedure was repeated twice.

#### Optic flow

To characterize responses to unpolarized motion stimuli, a sparse random dot pattern was presented on the LED display that simulated forward translational optic-flow (thrust), with the frontal point of expansion approximately at the eye-equator. Approximately 1% of LEDs in the display were illuminated in each frame of the pattern, with all other LEDs off (0.83 Weber contrast). Windowed regions of this pattern were presented sequentially (lateral-left: −90°:-50° azimuth; frontal: −40°:+40° azimuth; lateral-right: +50°:+90° azimuth; each covering the full elevation extent of ± 30°) followed by the whole pattern (−90°:+90° azimuth). Motion was presented in each region for 4 s, with an inter-trial period of 4 s with all LEDs off. This procedure was repeated twice.

### Data analysis

#### Data export

Recorded imaging data was exported as 8-bit tiff frames. We compiled all time-points for a single imaging plane and a maximum average intensity projection (MIP, detailed below) across all planes at each time-point.

#### Image registration

We used a DFT-based registration algorithm (Guizar-Sicairos et al., 2008) to first correct for motion in the MIP of the activity-independent tdTomato channel across all timepoints. We then applied the same registration displacements (*x,y*) to all individual planes of the activity-dependent GCaMP channel.

#### Maximum intensity projection

We constructed a maximum intensity projection (MIP) based on each imaging plane’s time-averaged fluorescence intensities, which avoided a bias towards including cells that were bright throughout an experiment but did not necessarily show modulation (versus cells which were inhibited for the majority of an experiment but were modulated nonetheless). The time-series of each pixel in the projection also originated from a fixed plane throughout the recording. In summary: for each imaging plane, we found an average intensity image sampling only frames captured during periods of inactivity between stimulus sets. We then found the imaging plane (*z*) with the highest average intensity at each position (*x,y*). The intensity time-series (*t*) from this location (*x,y,z*) was then inserted into a new array (*x,y,t*) to form the projection. Neighboring pixels in the projection could therefore contain signals from different imaging planes, but individual pixels contained signals from only one plane. All analysis was conducted on this projection unless otherwise stated.

#### Angle of polarization tuning

For each pixel, we found the average fluorescence intensity across the frames captured during each angle presentation to obtain a polarization tuning curve. Since a polarization-tuned analyser should respond identically to parallel angles of polarization (e.g. 0°/180°), we expected bimodal data with diametrically opposite modes. We therefore found the axial mean resultant vector, correcting for grouped data, and took its angle as the preferred angle of polarization, defined modulo 180° (Batschelet, 1965; Berens, 2009; Zar, 1999).

#### Polarization-selectivity index

For each pixel, we found the average fluorescence intensity during the first two presentations of the angles closest to and diametrically opposite its preferred angle of polarization in the tuning experiment (*F*_*pref*_). We then found the average intensity at orthogonal angles (*F*_*ortho*_) and calculated the polarization-selectivity index (PSI) as the difference between *F*_*pref*_ and *F*_*ortho*_, divided by their sum, with possible values ranging from 0 to 1. Where average PSI values are reported for a driver line, we used a broad ROI drawn around all labeled neurons in the brain area recorded, which we refer to as the ‘overall ROI’. To draw the overall ROI we used an average intensity image from frames between stimulus sets as a guide. We also used this average intensity image to define additional regions: we defined regions of ‘cells’ as the brightest 10% of pixels within the overall ROI, unless otherwise stated (e.g. Fig. 5B,C), and ‘background’ as the dimmest 10% of pixels outside of the overall ROI. For the overall ROI, cells and background regions, the distribution of PSI values within a recording tended to be non-normal; for average values we report the median value for an individual animal and the mean of the median values across animals. Where ΔPSI values are reported, we subtracted the mean PSI values within the same region across all tuning experiments recorded with the polarizer removed. Where we applied a PSI-threshold to filter polarization-selective pixels in a recording (e.g. tuning maps, polarotopy analysis), we used the mean + 1 SD of PSI values within its background. This typically resulted in a PSI threshold between 0.3–0.4. This threshold was modified for E-PG recordings in the protocerebral bridge where PSI values of cells tended to be lower than the background when averaged over multiple presentations; instead we used the mean + 1 SD of PSI values within cells across all tuning experiments with the polarizer removed.

#### Polarization tuning maps

To construct spatial maps of polarization tuning, we combined a color-coded representation of preferred angle of polarization and a grayscale representation of average intensity (Fig. S1J). Pixels falling within the overall ROI which had an above-threshold PSI value (see *Polarization-selectivity index*) were assigned a color consistent with those used previously (Weir et al., 2016) to convey their preferred angle of polarization. All other pixels with below-threshold PSI value or falling outside of the overall ROI convey their average intensity during periods of inactivity with a normalized grayscale color-code (Fig. S1J).

#### Automatically generated ROIs

In addition to manually drawn ROIs, we generated ROIs based on polarization tuning maps (Fig. S2A). Briefly, we discretized tuning maps so that they contained only 6 preferred angles of polarization, corresponding to those presented in the tuning experiment ± 15°, plus null values for excluded pixels. For each angle, we identified contiguous areas of 20 or more pixels with that tuning and retained the largest area as an ROI.

#### Time-series

We found the mean fluorescence intensity of pixels within a given ROI in each frame to obtain its time-series (F_*t*_). For polarization tuning experiments, we calculated ΔF/F = F_t_/F_0_-1, where F_0_ was the root mean square value of the time-varying intensity across the entire experiment. For all other experiments, we calculated F_0_ as the mean of F_t_ during the 0.5 s preceding stimulus onset. To find the average time-series across multiple recordings with mismatched sampling times, we resampled values at a common rate using linear interpolation. This procedure produced no discernible alteration of the original data points.

#### Polarotopy and scatter plots

For recordings in the medulla and AOTU, we included only the set of polarization-selective pixels, as described for the tuning maps (see *Polarization tuning map*). For recordings in the bulb and protocerebral bridge, we used ROIs drawn manually on individual glomeruli. We projected pixel or ROI positions (*x,y*) onto a single horizontal axis (anterior-posterior in the medulla, medial-lateral in the central brain) or vertical axis (ventral-dorsal throughout) and then normalized to give a linear position ranging from 0 to 1. The majority of recordings were performed in the right brain hemisphere; where left hemisphere recordings were included, we inverted their positions along both axes (i.e. in the medulla, anterior positions on the left were pooled with posterior positions on the right), since we expected the mirror-symmetric polarotopy found in the dorsal rim (Fig. S1G,H) to be preserved downstream. We then pooled the normalized positions and corresponding preferred AoP across all recordings and created a scatter plot with a random subset of 1000 data points, displaying either the corresponding PSI value or preferred AoP as the color of each point in the plot.

We quantified circular-linear associations between preferred angle (multiplied by two to correct for axial data) and normalized position by finding the slope and phase offset of a regression line, and then a correlation coefficient, according to (Kempter et al., 2012). We found the correlation coefficient for the population by pooling all data points, then performed a permutation test on the pooled dataset with shuffled combinations of position and preferred AoP and recalculated the correlation coefficient 10,000 times. We report an upper-bound on the p-value as the proportion of shuffled datasets with a correlation coefficient exceeding that found for the experimental dataset plus one (Phipson and Smyth, 2010). We also found the correlation coefficients for individual recordings and an associated p-value (Kempter et al., 2012). Where indicated, the regression lines for the pooled dataset and for individual recordings with a sufficient number of pixels to give a meaningful correlation (p<0.05) are shown on scatter plots.

We applied the Fisher z-transformation to correlation coefficients to find a mean correlation coefficient across flies. We used a hierarchical bootstrap method (Saravanan et al.) to find 95% confidence intervals for the mean correlation coefficient found. We resampled with replacement from the population of flies, then resampled with replacement from all recordings made from those flies and recalculated the mean correlation coefficient after applying the Fisher z-transformation, repeated 10,000 times. From the bootstrapped population of mean correlation coefficients we found confidence intervals using the bias-corrected and accelerated method (Efron, 1987). In all cases, the correlation coefficient for the pooled dataset from all recordings was found to be close to the mean coefficient for individual flies and within the confidence interval calculated. For recordings in the bulb and protocerebral bridge, we also calculated the circular-circular correlation coefficient (Berens, 2009; Zar, 1999).

#### Polar histograms

We found the normalized probability distribution of preferred angles of polarization with a bin width of 15°. We then constructed polar histograms with each bin’s probability depicted as the area of a wedge, rather than its radial length. We included in this analysis either all pixels within the overall ROI (Fig. 7) (see *Polarization-selectivity index*) or the region of cells only (Fig. 5) (see *Polarization tuning maps*), in which case we excluded recordings with few above-threshold pixels (less than 10% of the overall ROI). The results were qualitatively similar in both cases.

#### Population tuning vectors

For individual recordings, we found the direction and length of the population tuning in an individual animal by calculating the axial mean resultant vector of its preferred angles of polarization. For the pixel-based approach, we included all pixels within the overall ROI and weighted individual preferred angles by their PSI value (Berens, 2009), rather than applying a threshold. Since individual neurons with a larger area provided a greater contribution in this analysis we compared it with an ROI-based approach, using ROIs drawn manually on individual micro-glomeruli in the bulb. We excluded recordings with fewer than four ROIs, and weighted the individual preferred angle of an ROI by its mean PSI-value. The results were qualitatively similar for both approaches.

#### Cross-correlation

For E-PG recordings in the protocerebral bridge, we manually drew ROIs on the 16 individual glomeruli visible in each recording (one additional column on either end of the PB does not contain E-PGs). We then paired each ROI on the left side with an ROI on the right side, using a pairing scheme which wrapped on either side independently (i.e. 1L/1R, 8L/2R, 7L/3R, see Fig. 8A). For each pair, we obtained the time-series for the ROIs across all frames in the recording and found their normalized cross-correlation coefficient at zero lag, ranging from −1 to 1. We plot the coefficient values for each pair (Fig. S6A) and the mean coefficient across all pairs from all recordings after applying the Fisher z-transformation. We then shifted the pairing scheme by one position on the right side and repeated the procedure until all pairing schemes had been evaluated.

#### Auto-correlation

For recordings in the bulb, we used ROIs manually drawn on individual micro-glomeruli. For E-PG recordings in the protocerebral bridge, we used ROIs drawn on pairs of left and right glomeruli (Fig. 8A). For each ROI, we obtained the time-series across the first two cycles of the tuning experiment. We detrended the time-series and calculated its normalized auto-correlation function. We then found the time difference between the first peak in the function and the period of the stimulus presented during the tuning experiment. We plot the value of these time differences for each ROI, which we refer to as a ‘peak shift’ (Fig. S6D), along with limits for the maximum expected peak shift for a phase-locked response to the stimulus (± 2 s, half the duration of each angle presentation).

#### Data and code availability

The datasets and code generated during this study are available at the Open Science Framework: doi.org/10.17605/osf.io/3tsd6

### Confocal imaging

#### Fly lines

The following driver lines belonging to the Janelia (R) (Jenett et al., 2012) and Vienna Tiles (VT) (Tirian and Dickson, 2017) collections, were obtained from Bloomington Drosophila Stock Center (BDSC): R13E04-Gal4 (48565), R13E04-LexA (53457), R13E04-p65.AD (isolated from original stock number: 86690), VT059781-Gal4.DBD (75090), R56F07-Gal4 (39160), R73C04-Gal4 (39815), R17F12-Gal4 (48779), R49E09-Gal4 (38692), R88A06-Gal4 (46847), R34H10-Gal4 (49808), R34D03-Gal4 (49784), R34D03-LexA (54662), R19C08-Gal4 (48845), R78B06-Gal4 (48343).

The following stocks were also acquired from BDSC: Pan-R7-Gal4 (II; 8603), Pan-R7-Gal4 (III; 8604), 10xUAS-mCD8::GFP (32184), 26xLexAop-mCD8::GFP (32207), [10xUAS-mCD8::RFP, 13xLexAop-mCD8::GFP] (32229), UAS-sytGCaMP6s (64415), UAS-tdTomato (36328), MCFO-4 (64088), MCFO-5 (64089), MCFO-6 (64090), [UAS-nsyb-spGFP1-10, LexAop-CD4-spGFP11] (GRASP; BDSC 64314). *trans*-Tango (77123) was provided by G. Barnea. SS00096-Gal4 was a gift from V. Jayaraman and T. Wolff.

#### Fly rearing for immunostaining

Flies were raised at 25°C on a standard cornmeal/molasses diet in bottles or vials, under a 12:12 hour dark:light cycle, and we dissected 3–4 day old female flies. For *trans*-Tango analyses we dissected 17–18 day old female flies raised at 18°C (Talay et al., 2017).

#### Immunostaining

Immunohistochemical staining was conducted as previously described (Omoto et al., 2017; 2018). Briefly, brains were dissected in phosphate buffered saline (PBS) and fixed in ice-cold 4% EM-grade paraformaldehyde in PBS for 2.5 hours. They were subsequently washed for 4 × 15 mins in ice-cold PBS followed by cold ethanol dehydration (5 min washes in 5, 10, 20, 50, 70, 100% EtOH). After incubation for approximately 12 hours in 100% EtOH at 4°C, brains were subjected to a rehydration procedure with EtOH in the reverse sequence. Brains were then washed for 4 × 15 min in ice-cold PBS and 4 × 15 min in ice-cold 0.3% PBT (PBS with 0.3% Triton X-100), followed by 4 × 15 min in room temperature (RT) 0.3% PBT. They were then incubated in blocking buffer (10% Normal Goat Serum in 0.3% PBT) for 30 min at RT. Following this, the brains were incubated in primary antibodies, diluted in blocking buffer at 4°C for approximately three days. They were subsequently washed 4 × 15 min in RT 0.3% PBT and placed in secondary antibodies diluted in blocking buffer at 4°C for approximately three days. They were finally washed 4 × 15 min in RT 0.3% PBT and placed in VectaShield at 4 °C overnight before imaging (Vector Laboratories). *trans*-Tango and GRASP analyses required separate staining of neuropil and respective fluorophores due to different incubation times.

The following antibodies were used: rat-antiDN-cadherin (DN-EX #8, 1:20, Developmental Studies Hybridoma Bank); mouse anti-neuroglian (BP104, 1:30, Developmental Studies Hybridoma Bank); chicken anti-GFP (1:1000, ab13970, Abcam); Rabbit anti-DsRed (1:1000, 632496, Clontech); rabbit anti-HA (1:300, Cell Signaling Technologies); and mouse anti-V5 (1:1000, ThermoFisher Scientific). The following secondary antibodies, IgG_1_ (Jackson ImmunoResearch; Molecular Probes, Thermo Fisher Scientific), were used: Cy5 conjugated anti-mouse (1:300), Cy3-conjugated anti-rat (1:300), Alexa 488-conjugated rabbit-anti-GFP (1:1000), Alexa 488-conjugated anti-chicken (1:1000), Alexa 546-conjugated anti-rabbit (1:1000), and Alexa 488-conjugated anti-mouse (1:1000). The following antibodies from Abcam were also used: Cy5-conjugated anti-rat (1:300) and Cy3-conjugated anti-rabbit (1:300).

#### Confocal microscopy and image analysis

Processed brains were mounted on glass slides and imaged in either the antero-posterior (A–P) or dorsal-ventral (D–V) axis with a Zeiss LSM 700 Imager M2 using Zen 2009 (Carl Zeiss), with a 40x oil objective. Images were processed using Image J (FIJI) (Schindelin et al., 2012). Image stacks of the AOTU or EB were rotated slightly and interpolated to align the neuropil with the imaging plane. Background labeling was removed to improve visualization in some projections (Fig. 2B,C, Fig. 3G–G’’’).

## SUPPLEMENTARY INFORMATION

**Figure S1:**
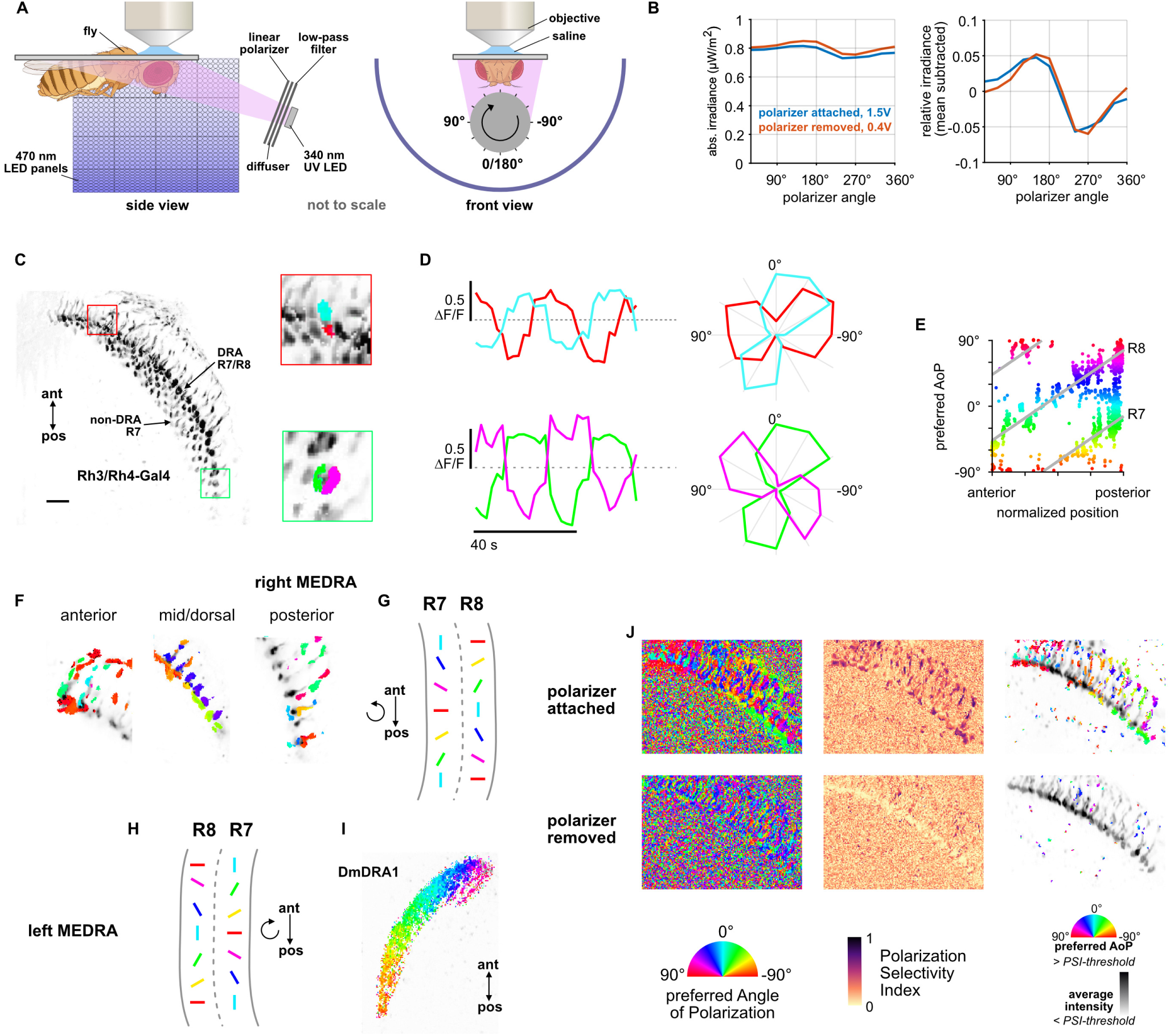
Polarizer characterization and R7/R8 photoreceptor stimulation. **A**: Schematic of experimental setup. Volumetric two-photon imaging of the medulla dorsal rim area (MEDRA) was performed while ultraviolet light was presented continuously and a linear polarizing filter varied the angle of polarization. Rotations and angles of polarization are expressed from the external viewpoint looking towards the animal’s head. (Fly illustration: BioRender.com) **B**: Modulation of intensity over one revolution of the polarizer in absolute units (left) and with the mean subtracted (right). The amplitude of modulation (approximately ± 5%) was similar with the polarizer attached or removed. **C**: Example time-averaged maximum-intensity projection of GCaMP activity in DRA R7/R8 + non-DRA R7 photoreceptors in the dorsal medulla (Rh3/Rh4-Gal4>sytGCaMP6s). Insets: ROIs drawn on R7 and R8 terminals in anterior (top) and posterior (bottom) MEDRA. **D**: GCaMP activity in R7/R8 terminals from **C** in response to rotations of polarizer. Right: Polar plot of average responses for each angle of polarization presented. **E**: Example scatter plot showing the polarotopic organization of DRA R7/R8 photoreceptors for the recording in **C**. Individual points represent pixels recorded from R7/R8, showing their normalized horizontal position in the MEDRA and their preferred angle of polarization (AoP). **F**: Example tuning maps of preferred AoP for recordings in a single plane, showing details of R7/R8 terminals in posterior, mid/dorsal and anterior MEDRA in the right optic lobe. **G**: Summary of preferred AoP in R7/R8 in the right MEDRA (from Weir et al., 2016). **H**: Summary of preferred AoP in R7/R8 in the left MEDRA. **I**: Example polarization tuning map for DmDRA1 in the left MEDRA. **J**: Example construction of a polarization tuning map for a maximum-intensity projection of two-photon imaging data in the medulla. Left: Preferred AoP for all pixels, with the polarizer attached (top) and removed (bottom). GCaMP-expressing photoreceptors can be differentiated from background noise, and show a retinotopic organization of preferred AoP only with the polarizer attached. Center: Polarization-selectivity index (PSI), a measure of fluorescence intensity modulation by the polarizer device, for the same data. Right: Preferred AoP values with a PSI-threshold applied. Below-threshold pixels (grayscale) show average intensity values over the experiment.

**Figure S2:**
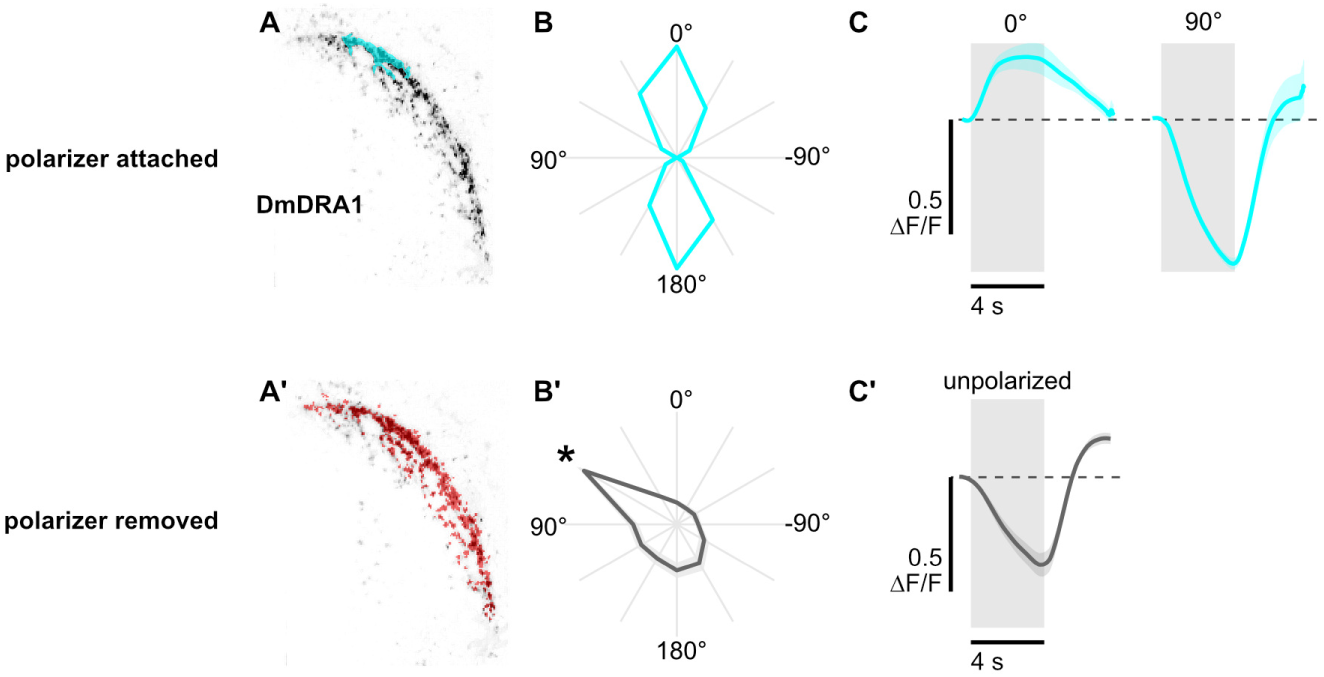
Polarization-opponent flash responses in DmDRA1. **A**: Example time-averaged maximum-intensity projection showing GCaMP activity in DmDRA1 neurons (DmDRA1-split>sytGCaMP6s) and example ROIs automatically-generated around areas of DmDRA1 neurons with a preferred angle of polarization around 0° (top, cyan) or around the brightest pixels for experiments with the polarizer removed (bottom, red). **B**: Normalized tuning curves for ROIs (N = 11, one ROI per animal). Mean ± SEM. **B**’: *denotes the first angle of polarization presented, during which time activity was often falling in experiments with the polarizer removed (see Fig. 1C). **C**: Average responses of ROIs to 4 s UV light flashes with the polarizer at 0° (pk ΔF/F = 0.23) and 90° (pk ΔF/F = −0.64, N = 10, p = 0.0002), and with the polarizer removed (bottom) (pk ΔF/F = −0.38, N = 7). Mean ± SEM.

**Figure S3:**
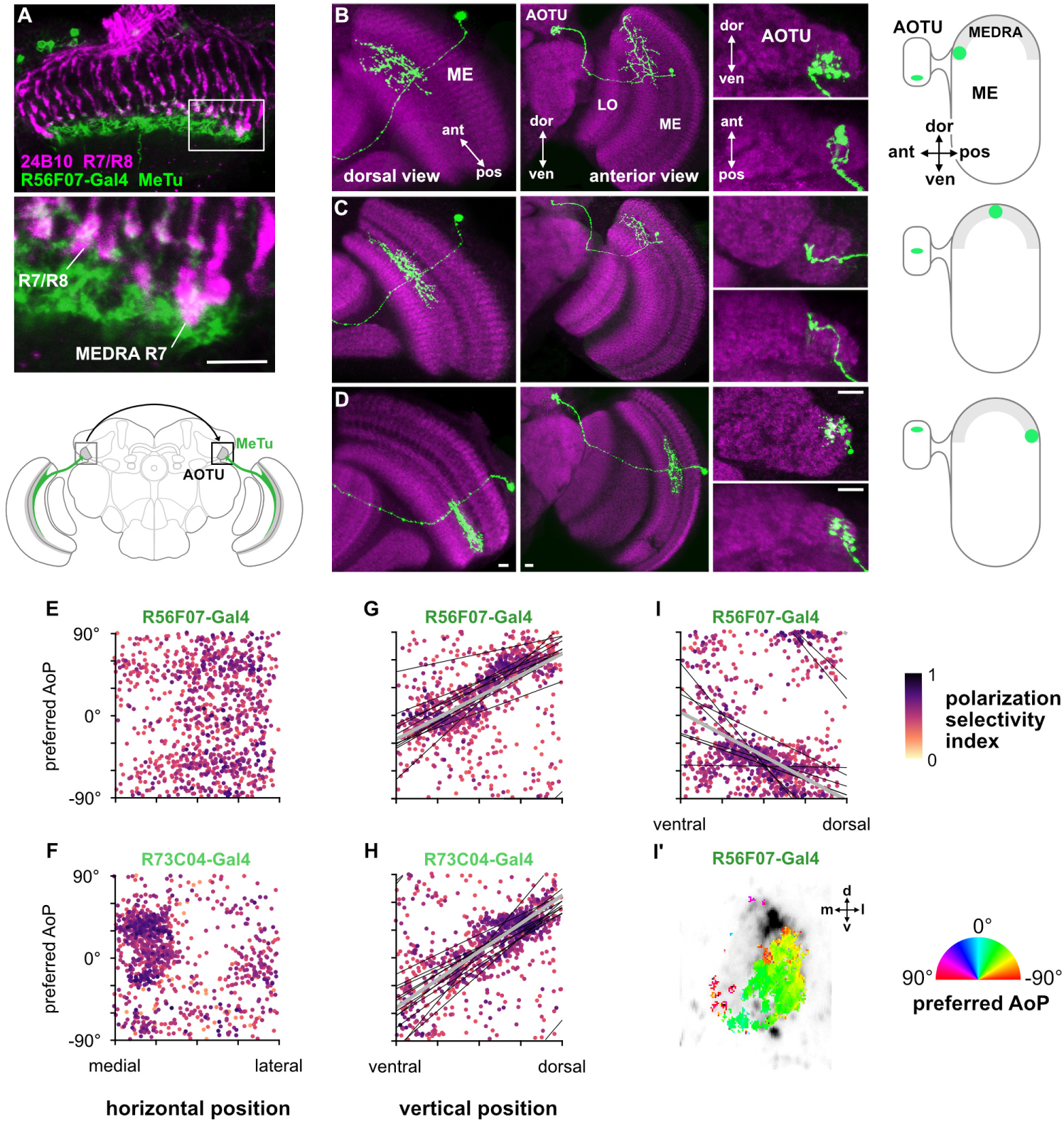
Retinotopic mapping of medulla dorsal rim area to AOTU by MeTu neurons and organization of polarization-selective responses. **A**: Confocal section of the medulla (dorsal view) showing R7/R8 photoreceptors (24B10 antibody staining: green) and their proximity to MeTu neurons (R56F07-Gal4>GFP: magenta). Bottom: Enlargement of medulla dorsal rim area (MEDRA). Scale bar denotes 10 μm. **B**: Confocal projections of a single MCFO clone of R56F07 MeTu neurons with dendrites in the anterior/dorsal medulla (ME) in proximity to the medulla dorsal rim area. Left: Dorsal view. Center: Anterior view. Right: High magnification projections showing the position of terminals in the anterior optic tubercle (AOTU). **C**: As in **B**, for a MeTu neuron with dendrites in the mid/dorsal medulla. **D**: As in **B**, for a MeTu neuron with dendrites in the posterior/dorsal medulla. Scale bars denote 10 μm. **E**: Scatter plot showing the organization of polarized light responses in R56F07 MeTu neurons. Individual points represent pixels recorded in MeTu neurons, showing their normalized horizontal position in the AOTU and their preferred angle of polarization (AoP). Color displays PSI value (pooled ρ = 0.03, N = 17 recordings). **F**: As in **E**, for R73C04 MeTu neurons (pooled ρ = −0.22, N = 11 recordings). **G**: Scatter plot showing the predominant polarotopic organization of R56F07 MeTu neurons. Thin lines show linear-circular fits for data from individual animals with significant correlations (mean individual ρ = 0.61, SEM 0.16, N = 7 animals), thick line shows fit for all pooled data (pooled ρ = 0.68, N = 8 recordings, p < 10^−6^ permutation test). **H**: As in **G** for R73C04 MeTu neurons (mean individual ρ = 0.68, SEM 0.12, N = 10 animals), thick line shows fit for all pooled data (pooled ρ = 0.58, N = 10 recordings, p < 10^−6^ permutation test). **I**: Scatter plot showing an occasional, second organization of responses in R56F07 MeTu neurons (mean individual ρ = 0.52, SEM 0.12, N = 6 animals), thick line shows fit for all pooled data (pooled ρ = 0.30, N = 7 recordings, p < 10^−6^ permutation test). **I’**: Example polarization tuning map of second organization of responses.

**Figure S4:**
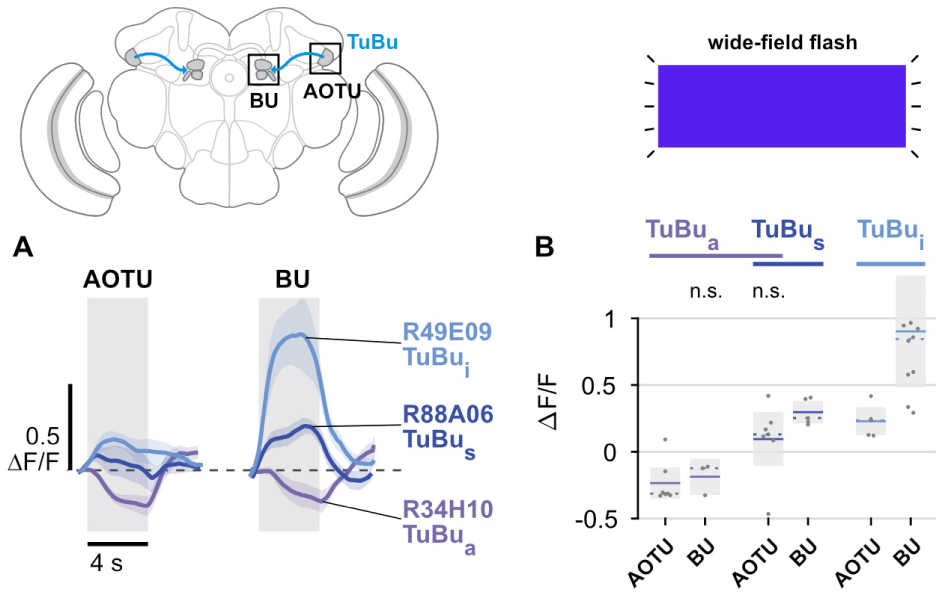
Unpolarized flash responses in TuBu neurons. **A**: Average responses of all TuBu neurons in each population to 4 s blue light flashes, recorded in the anterior optic tubercle (AOTU) (GCaMP6s) and bulb (BU) (sytGCaMP6s). Mean ± SEM. **B**: Peak responses for individual animals and their mean and median (dashed line). (pk ΔF/F **TuBu**_**a**_ AOTU: −0.23, CI 0.16, N = 7, p = 0.008, BU: −0.19, CI 0.12, N = 3, p = 0.11; **TuBu**_**s**_ **+ TuBu**_**a**_ AOTU: 0.10, CI 0.27, N = 7, p = 0.38, BU: 0.30, CI 0.10, N = 5, p = 0.02; **TuBu**_**i**_ AOTU: 0.23, CI 0.12, N = 5, p = 0.013, BU: 0.90, CI 0.68, N = 10, p = 0.002) Shaded box denotes Bonferroni corrected 95% confidence interval.

**Figure S5:**
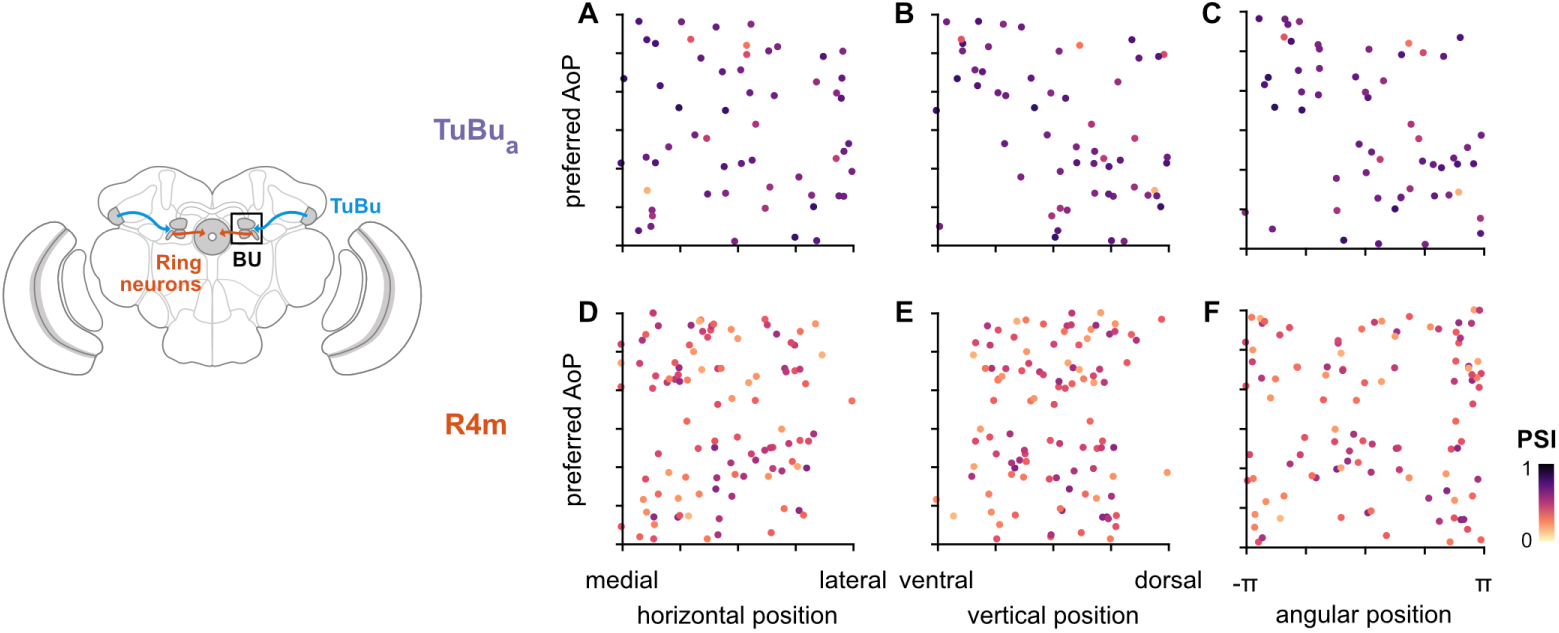
Unstructured organization of preferred angles of polarized light in the anterior bulb. **A**: Scatter plot showing the horizontal organization of TuBu_a_ tunings in the anterior bulb (BUa). Individual points represent ROIs drawn on micro-glomeruli, showing their normalized horizontal position within the BUa and their preferred angle of polarization (AoP). Color of individual points displays PSI value (**TuBu**_**a**_: N = 8 animals, 14 recordings, 6 left BU: 29 ROIs, 4.8 ± 1.0 per animal, 8 right BU: 28 ROIs, 4.7 ± 0.8 per animal; mean ROI PSI 0.65 ± 0.12) (0 significant individual linear-circular correlations; pooled data ρ = −0.02, p = 0.91 permutation test). **B**: As in **A**, for vertical organization of TuBu_a_ tunings (1 significant individual linear-circular correlation, ρ = −0.61; pooled data ρ = 0.46, p = 0.002 permutation test). **C**: As in **A**, for circular organization of TuBu_a_ tunings (5 significant individual circular-circular correlations, mean ρ = 0.84, SEM 0.69; pooled data ρ = −0.43, p = 0.23 permutation test). **D**: As in **A**, for horizontal organization of R4m tunings (**R4m**: N = 25 animals, 26 recordings, 2 left BU: 8 ROIs, 4.0 ± 0.0 per animal, 24 right BU: 96 ROIs, 4.0 ± 0.8 per animal; mean ROI PSI 0.38 ± 0.12) (1 significant individual linear-circular correlation, ρ = −0.76; pooled data ρ = 0.01, p = 0.96 permutation test). **E**: As in **B**, for vertical organization of R4m tunings (0 significant individual linear-circular correlations; pooled data ρ = 0.09, p = 0.47 permutation test). **F**: As in **C**, for circular organization of R4m tunings (3 significant individual circular-circular correlations, mean ρ = 0.98, SEM 0.34; pooled data ρ = 0.02, p = 0.98 permutation test).

**Figure S6:**
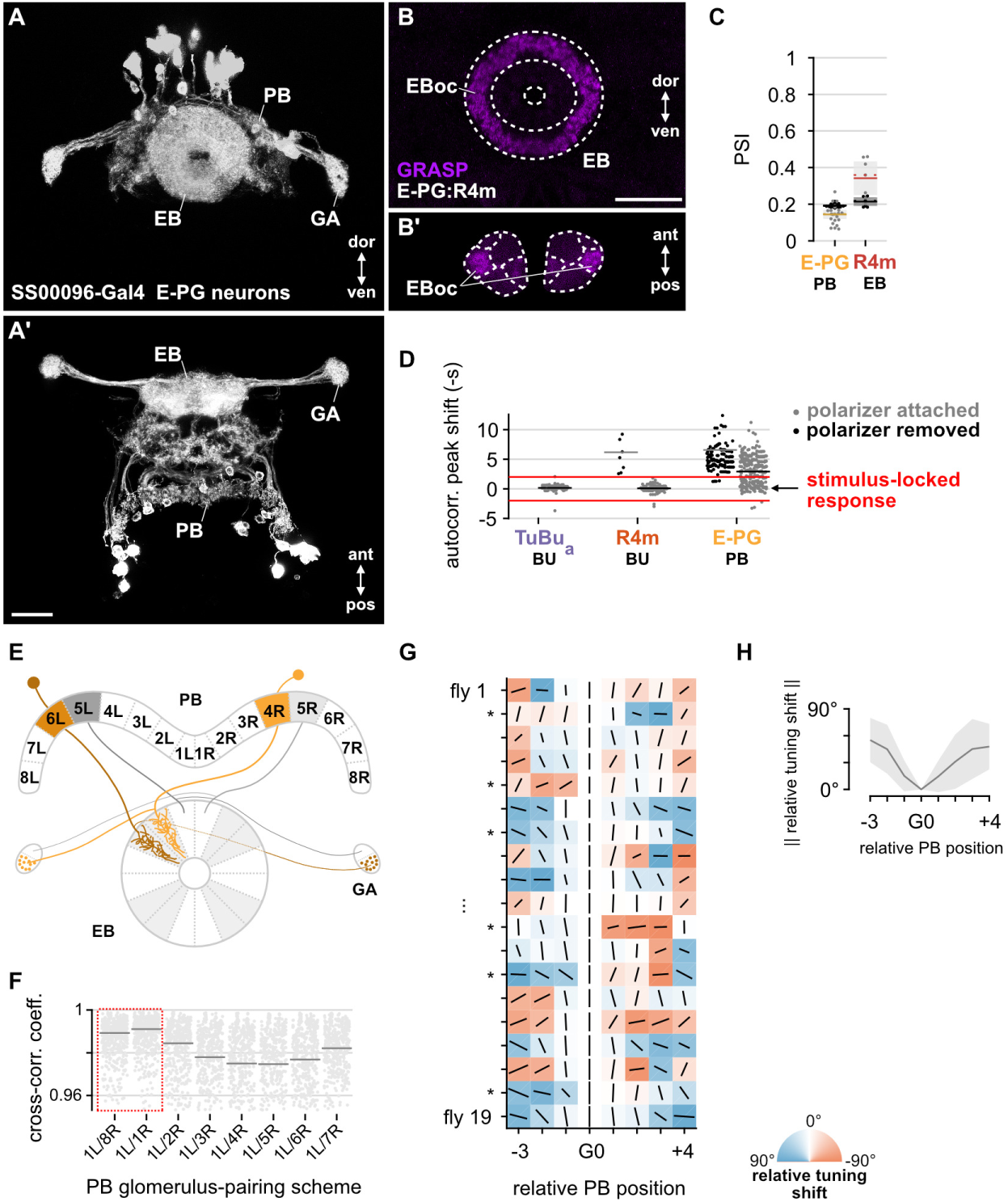
E-PG neurons show inconsistent responses to the angle of polarized light and variable tunings. **A**: Confocal projection (anterior view) of E-PG expression pattern in the ellipsoid body (EB), protocerebral bridge (PB) and gall (GA) (SS00096-Gal4>GFP). **A’**: Dorsal view. Scale bar denotes 25 μm. **B**: Confocal projection of GRASP (GFP reconstitution across synaptic partners) signal for connections from E-PG to R4m neurons in the EB. **B’**: Dorsal view. Scale bar denotes 25 μm. **C**: Average PSI values within E-PG neurons in the PB and R4m neurons in the EB (light dots) and background regions (dark dots) in individual animals (**E-PG** neurons: 0.14, CI 0.05, background: 0.19, CI 0.01, N = 22 animals, p = 0.0001 t-test; **R4m** neurons: 0.34, CI 0.11, background: 0.21, CI 0.03, N = 7 animals, p = 0.02 t-test). **D**: Shift in time of the first peak of an ROI’s auto-correlation function, relative to the period of the polarizer (0 s). Red lines indicate a window of ± 2 s: a peak shift of greater magnitude indicates a response which was not phase-locked with the polarizer stimulus (median peak shift **TuBu**_**s**_: attached 0.15 s, CI 0.59, N = 7 animals, 85 ROIs included; **R4m**: attached 0.07 s, CI 0.56, N = 25 animals, 126 ROIs included; removed 5.76 s, CI 8.91, N = 9 animals, 10 ROIs included; **E-PG**: attached 2.73 s, CI 2.77, N = 22 animals, 504 ROIs included; removed 4.79 s, CI 5.63, N = 18 animals, 175 ROIs included). **E**: Summary schematic of E-PG neuron innervation patterns in the ellipsoid body (EB) and protocerebral bridge (PB) and gall (GA). Highlighting indicates the L/R pairing scheme used. 9L/9R in PB not shown. **F**: Normalized cross-correlation coefficient for all E-PG pairs of left and right glomeruli in the PB, using different pairing schemes. Each scheme name gives the pairing of 1L and its right PB partner; all other pairs within the scheme follow the same logic. Horizontal lines mark the Fisher z-transformed mean coefficient (N = 22 animals). Highlighted schemes represent pairings of E-PGs innervating neighboring wedges of the EB. Pairing scheme 1L/1R is used in this study. **G**: Relative tunings in individual animals. Orientation of lines represent preferred AoP (relative to G0), length of lines indicate PSI (height of each square is equal to a PSI value of 1). Asterisks indicate significant individual circular-circular correlations between position and preferred AoP. **H**: Average tuning shift (relative to G0), summarizing data in **G**. Mean ± SEM (N = 19).

